# A high-quality genomic catalog of the human oral microbiome broadens its phylogeny and clinical insights

**DOI:** 10.1101/2025.03.10.642329

**Authors:** Jun Hyung Cha, Nayeon Kim, Junyeong Ma, Sungho Lee, Geon Koh, Sunmo Yang, Samuel Beck, Iksu Byeon, Byunguk Lee, Insuk Lee

## Abstract

Understanding the taxonomic and functional diversity of the human oral microbiome requires comprehensive genomic catalogs. We present the human reference oral microbiome (HROM), with 72,641 high-quality genomes from 3,426 species, including 2,019 novel species, significantly improving metagenomic sequence read classification over existing catalogs. Notably, HROM unveils 1,137 novel candidate phyla radiation (CPR) species, establishing Patescibacteria as the most prevalent phylum in the oral microbiota, with oral Patescibacteria forming a distinct clade from environmental Patescibacteria. We also identify an oral CPR subclade associated with periodontitis, which complement *Porphyromonas gingivalis* in predicting the disease. Finally, comparing HROM with reference genomes of the human gut microbiome reveals significant taxonomic and functional divergence between the oral and gut microbiomes. We identify 42 ectopic oral species and demonstrate that their relative abundance in gut microbiota is predictive of intestinal, cardiovascular, and liver diseases, highlighting the clinical importance of the oral microbiota in systemic disorders.

## Introduction

The human oral cavity serves as a gateway for microbial colonization. The microbial community within this environment is both structured and dynamic, with its collective functions having a significant impact on overall human health^1^. Dysbiosis of the oral microbiome has been linked to oral diseases such as dental caries and periodontitis^2^. Moreover, recent studies suggest a connection between oral microbiota and various systemic diseases, including gastrointestinal, cardiovascular, neurological, autoimmune diseases, and cancers^3^. As a result, understanding the diversity and functions of human oral microbiota is becoming increasingly important in healthcare research.

The initial survey of the human microbiome revealed that the gut and oral cavity are two primary body sites with highly diverse microbial communities^4^. Recent advances in genome-resolved metagenomics have uncovered numerous novel species through metagenome-assembled genomes (MAGs), a culture-independent method that reveals the true diversity of the biome and identifies bacterial genomes previously undetectable using culture-based approaches^5^. In the case of the human gut microbiome, several studies have demonstrated that the MAG strategy, involving sequential steps such as assembly and binning, has successfully identified under-represented clades and expanded our ecological and functional understanding of the previously unknown repertoire of the gut microbiome^6–8^. However, efforts to catalog the oral microbiome using MAGs have been limited in comprehensiveness and quality^9^. The most widely used genome database for the human oral microbiome, eHOMD^10^, primarily provides genomes for oral prokaryotic species, which are predominantly based on isolate genomes.

We have assembled MAGs from publicly available whole metagenomic sequencing (WMS) data from 7,956 diverse oral samples, alongside isolate genomes from the human oral cavity, resulting in the Human Reference Oral Microbiome (HROM). This genomic catalog provides 145,149 prokaryotic genomes of medium quality (MQ, completeness ≥ 50%, contamination < 5%), with 72,641 near-complete genomes (NC, completeness ≥ 90%, contamination < 5%) comprising 3,426 species. The HROM expands the phylogeny of the oral microbiome, including numerous novel species within the phylum Patescibacteria, a candidate phyla radiation (CPR) group^11^ known for its enigmatic lifestyle. Our analysis revealed that oral Patescibacteria species form a distinct clade compared to those found in the environment, with a subclade associated with the oral disease periodontitis. Additionally, the HROM facilitated a comparison between oral and gut microbiota, uncovering significant differences in their taxonomic and functional landscapes. Notably, we identified 42 oral bacterial species that are ectopically located in the gut, offering novel clinical insights into systemic diseases. With its extensive and high-quality genomic information, the HROM opens new avenues for innovative oral microbiome research, poised to reveal new dimensions of oral microbial ecology and its impact on human health.

## Results

### Cataloging oral microbial genomes using clade-adjusted quality evaluation methods

For cataloging microbial genomes in the human oral environment, we collected WMS data from 7,956 oral samples from publicly available resources such as ENA^12^, SRA^13^, and CNGBdb^14^ (downloaded in May 2022, **Supplementary Table 1**). These data represent samples from 21 countries, with the majority from the United States and China **(Fig. 1a)**. The samples cover various biogeographic sources within the oral cavity, with saliva and dental samples being the most common **(Fig. 1b)**. Most of WMS data from oral samples show high contamination rate of human DNA sequence reads (**Extended Data Fig. 1a**). Using a bioinformatic pipeline previously employed for cataloging gut microbial genomes^7^, we constructed MAGs **(Fig. 1c, Extended Data Fig. 1b)**. Additionally, we manually searched for prokaryotic genomes from the human oral cavity in public resources such as eHOMD (v.3)^10^, GenBank^15^, RefSeq^16^, PATRIC^17^, and a pre-existing catalog^18^ **(Supplementary Table 2)**.

**Fig. 1.**
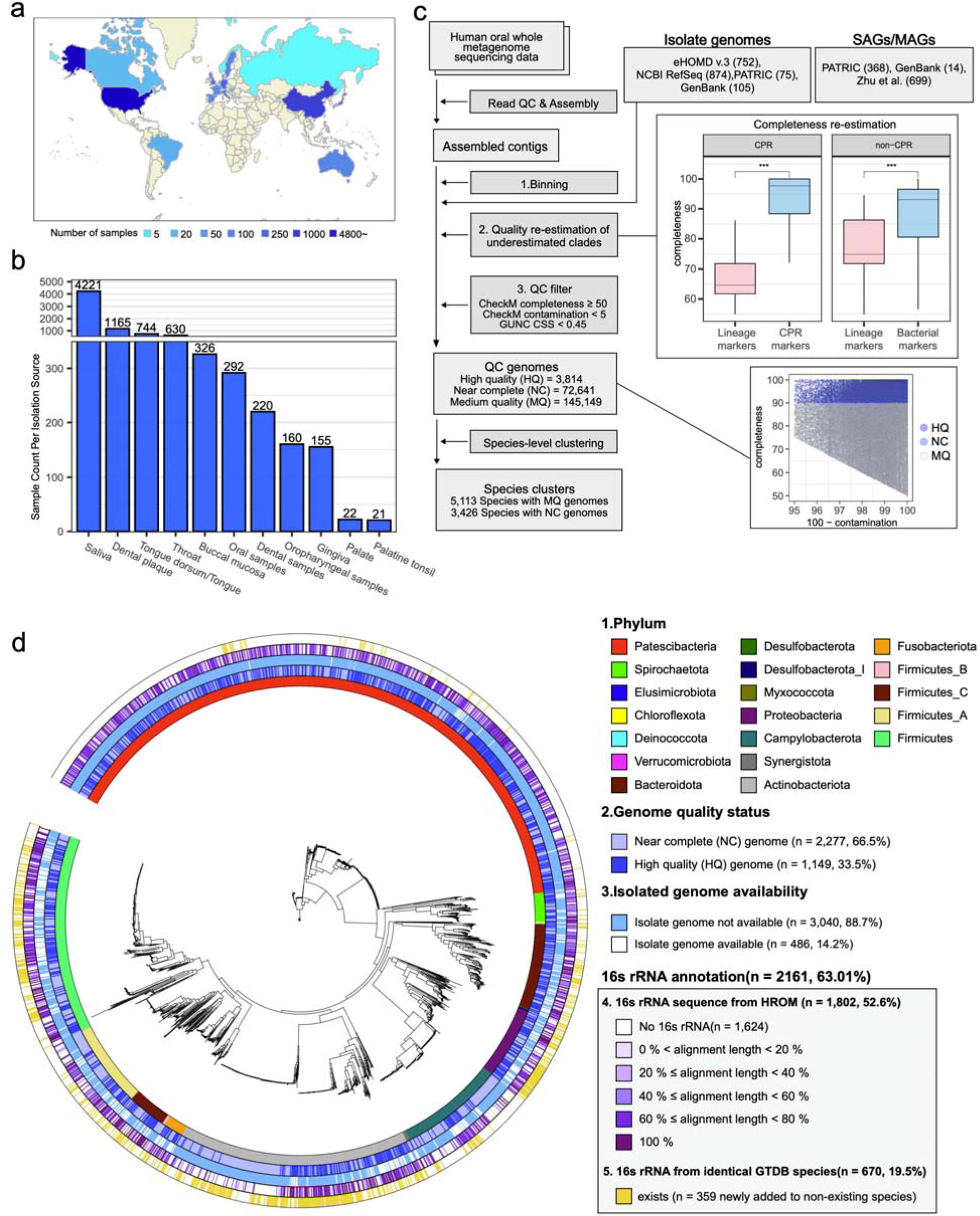
Overview of the human reference oral microbiome (HROM) construction and phylogenetic diversity. **a**, Geographic distribution of 7,956 whole metagenome sequencing (WMS) datasets from human oral cavities, representing samples collected across various countries. **b**, Distribution of WMS datasets across diverse isolation sources within the oral cavity, highlighting the number of samples for each source. **c**, Bioinformatics workflow for constructing the HROM catalog, including read quality control (QC), assembly, genome binning, re-evaluation of completeness and contamination, and species-level clustering. Completeness differences in re-evaluated genomes are shown, with statistical significance assessed using a two-tailed Mann-Whitney U test (\*\*\**P* < 0.001). Boxplot elements: center line (median), box edges (25th and 75th percentiles), whiskers (1.5× interquartile range). **d**, Phylogenetic tree of 3,424 near-complete, species-level bacterial genomes in HROM. Tracks around the tree indicate (1) phylum-level classification, (2) genome quality status, (3) isolate genome availability, and (4) completeness of 16S rRNA annotations. Genome quality categories are defined in the workflow, and 16S rRNA annotations include genomes from HROM and external references.

Genome quality is evaluated based on completeness and contamination using CheckM^19^, which relies on lineage-specific markers. We found that the CheckM lineage module often underestimates completeness for relatively novel clades, as their lineage-specific marker genes are not well established. To address this, we identified clades with underestimated completeness by comparing them with universal bacterial marker genes^19^ and re-estimated their genome quality (**Supplementary Table 3**). For the phylum Patescibacteria, which lacks many universal bacterial marker genes, CPR marker genes^20^ were used for re-estimation. This quality re-estimation process significantly improved completeness of 12,606 genomes **(Fig. 1c, Extended Data Fig. 1c)**. The re-estimated completeness was further validated by showing high correlation with both the percentage of hits from GTDB 120 single markers, an independent marker gene set, and the completeness estimated by CheckM2^21^, an updated method that resolves the limitation of lineage-specific markers using a machine learning approach **(Extended Data Fig. 1d)**.

Additionally, genomes were subjected to quality control based on genome chimerism using the Genome UNClutterer (GUNC)^22^, excluding genomes with clade separation scores (CSS) exceeding 0.45. This resulted in a total of 145,149 non-redundant genomes meeting the MQ genome criteria, represented by 5,113 species clusters **(Supplementary Table 4-5)**. Of these genomes, 72,641 NC genomes represent 3,426 species, with 2,019 (59%) unclassified at the species level by GTDB-tk^23^, indicating their novelty. Species clusters consisting exclusively of MQ genomes typically contain only a single low-quality genome, making it unlikely that they correspond to real organisms. For example, clustering with MQ genomes yields 1,687 additional species; however, 1,374 of them (81.4%) are singletons. Furthermore, MQ genomes, which lack substantial portions of genomic content, can mislead functional analyses of species. Therefore, we focused on 3,426 species composed of NC genomes for HROM-based taxonomic profiling of microbiomes (**Fig. 1d**). Notably, only 486 of the 3,426 species (∼14.2%) have corresponding isolate genomes, and just 319 species (∼9.3%) include genomes from eHOMD, indicating that the majority of oral microbes remain uncultivated.

### HROM outperforms pre-existing genomic catalogs in oral WMS read classification

We compared HROM to pre-existing genomic catalogs for the human oral microbiome. Notably, the expanded human oral microbiome database (eHOMD)^10^, one of the most widely used human oral microbiome databases, predominantly consists of isolated/culturable genomes. Although eHOMD is frequently used as a reference genome database for oral microbiome analysis, it includes genomes not isolated from the human oral cavity. For instance, genome SEQF2350 (GenBank accession = GCA_000259485.1, biosample accession = SAMN00761858) is recorded as being human urogenital tract isolate. To address this issue, we manually inspected the genomes in eHOMD and selected only 752 confirmed oral microbial genomes from the 2,213 total genomes for inclusion in HROM, ensuring it comprises only authentic human oral-origin genomes.

HROM consists of 72,641 NC genomes, approximately 32.8 times the total number of genomes in eHOMD (**Fig. 2a**). When applying the same species-level clustering scheme, HROM included 7.37 times more species based on NC genomes (NC species) compared to eHOMD (**Fig. 2b**). We further compared HROM with other genome catalogs, including COGR^24^, human-oral-v1-0 microbiome catalog in MGnify repository^25^, and the largest existing MAG-based oral genome catalog of Zhu et al^9^, although the latter provides only representative genomes to the public. When applying the same genome quality criteria as HROM, only 1,048 NC species were identified in the Zhu et al. database. Overall, our analysis demonstrated that HROM contains substantially more NC genomes and corresponding species compared to other genome catalogs included in this comparison.

**Fig. 2.**
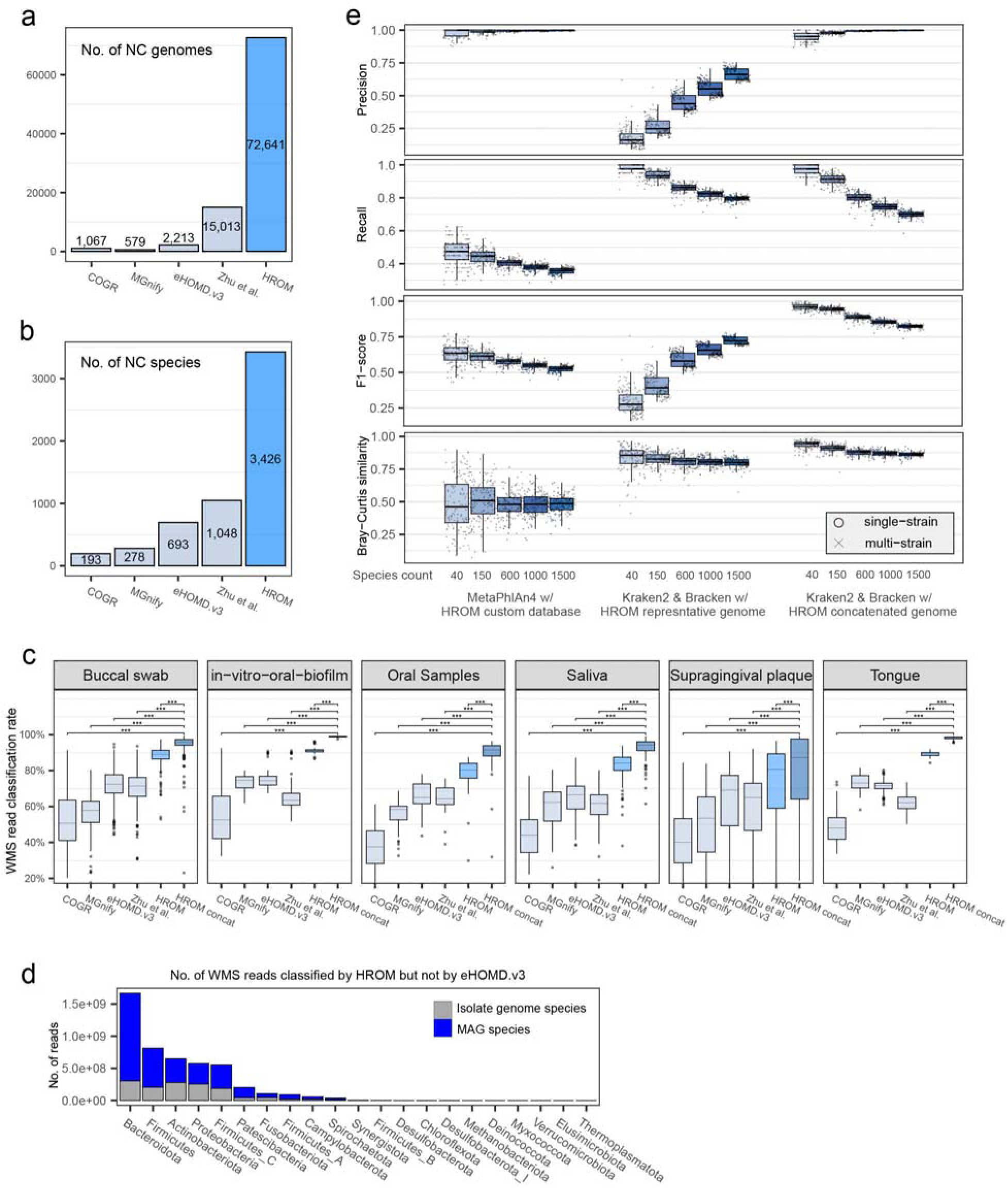
Comparative analysis and Benchmarking of the HROM catalog. **a**, Total number of near-complete (NC) genomes in the HROM catalog compared with other oral genome catalogs, including COGR, MGnify, eHOMD v3, and Zhu et al. For the Zhu et al. catalog, only representative genomes were publicly available, therefore, the total genome count is based on the reported statistics. **b**, Number of NC species identified in the HROM catalog compared to other genome catalogs. For Zhu et al., only representative genomes were publicly available; thus, we present the count of non-chimeric, NC representative genomes. **c**, Classification rates of WMS reads for different oral sample types (buccal swab, in-vitro oral biofilm, oral samples, saliva, supragingival plaque, and tongue) using various genome catalogs. For consistency, only non-chimeric and NC genomes were used for classification. Except for ’HROM concat,’ which represents the concatenated HROM database, only representative genomes were utilized. HROM achieves the highest classification rates across all sample types. Statistical significance was assessed using a two-tailed Mann-Whitney U test (*** *P* < 0.001). **d**, Total number of WMS reads from a validation dataset (PRJNA917836, buccal mucosa) that were classified by HROM but not by eHOMD v3, broken down by isolate genome species and MAG species, across different taxonomic phyla. **e**, Benchmarking performance of species-level taxonomic profiling methods using simulated datasets of varying species complexity (40, 150, 600, 1,000, 1,500 species). Metrics include precision, recall, F1-score, and Bray-Curtis similarity. Results are shown for MetaPhlAn4 with the HROM custom database, and Kraken2/Bracken with both HROM representative and concatenated genome databases, under single-strain and multi-strain scenarios.

The improved comprehensiveness of genome catalogs may also enhance the taxonomic classification of WMS reads. We aligned sequence reads from six independent public human oral WMS datasets, comprising 1,129 samples not used in the construction of HROM **(Supplementary Table 6)** to various genome catalogs using Kraken2^26^. HROM significantly outperformed other genome catalogs in read classification rate **(Fig. 2c)**. Furthermore, using concatenated conspecific non-redundant genomes for each HROM species further improved performance compared to using representative genomes alone. Sequence reads unaligned by eHOMD were primarily classified into the phyla Bacteroidota, Proteobacteria, Firmicutes, and Patescibacteria across most oral cavity isolation sources **(Fig. 2d, Extended Data Fig. 2a)**. These findings suggest that the enhanced oral WMS read classification capabilities of HROM will enable more accurate taxonomic profiling of human oral microbiomes.

### DNA-based species profiling with HROM outperforms marker-based profiling of the human oral microbiota

One of the major applications of genomic catalogs of microbiomes is assembly-free taxonomic profiling of WMS samples. Although a marker-based method, MetaPhlAn4^27^, is widely used, we expect that the improved read classification rate provided by HROM will enable more accurate species-level taxonomic profiles through DNA-based methods, such as alignment using Kraken2^26^ followed by reassignment using Bracken^28^. To compare these two approaches of species profiling of oral microbiome, we generated custom databases for both MetaPhlAn4 and Kraken2 based on HROM. For Kraken2, we created two versions of custom databases: one based on species representative genomes and the other on concatenated conspecific genomes for each species. While building Kraken2 custom databases from a given genome catalog is relatively straightforward, creating a custom database of species-specific markers for MetaPhlan4 requires significant efforts. This may explain why most pre-existing genomic catalogs provide custom databases for Kraken2 but not for MetaPhlan4. Based on the methodological principles of the MetaPhlAn4 protocols, we constructed a species-specific marker database for HROM, but marker genes could only be identified for 1,603 species (46.8% of HROM species; **Extended Data Fig. 2b**). We observed that species-specific marker genes could not be identified for species undergoing rapid speciation, characterized by widely dispersed branches in the phylogenetic tree, such as those in the phyla Patescibacteria and Actinobacteriota. These rapidly evolving species exhibit a high density of single nucleotide variations (SNVs)^7^ but lack gene-level diversification. The absence of marker genes for these clades may reduce the accuracy of HROM-based species profiling of the oral microbiome when using MetaPhlAn4.

The limited marker genes for fast-evolving clades may reduce the performance of species profiling using the MetaPhlAn4 compared to Kraken2. To evaluate the accuracy of these methods under varying species complexity, we generated simulated WMS samples using HROM genomes for communities containing 40, 150, 600, 1,000, and 1,500 species, with either single or multiple strains. Using WMS samples from healthy individuals, we surveyed the number of HROM species across oral cavity biogeographic locations. Except for the throat, most environments harbored between 700 and 1,100 species per individual (**Extended Data Fig. 2c**). To improve accuracy, we applied a Kraken confidence threshold of 0.2 to reduce false positives and performed genome-size normalization for better abundance measurement^29^. Both MetaPhlAn4 and Kraken2/Bracken with concatenated genomes achieved near-perfect precision in species profiling. However, MetaPhlAn4 showed much lower recall than Kraken2/Bracken. Consequently, the F1 score, which balances precision and recall, indicated that Kraken2/Bracken with the HROM custom database based on concatenated genomes performed best (**Fig. 2e**). Furthermore, Kraken2/Bracken outperformed MetaPhlAn4 in Bray-Curtis similarity-based evaluations of species abundance profiles. Finally, we confirmed that the application of the Kraken confidence threshold and genome-size normalization significantly improved species profiling accuracy (**Extended Data Fig. 2d**). These results demonstrate that the Kraken2/Bracken pipeline using concatenated species genomes, genome-size normalization, and a confidence threshold provides the most accurate species profiles for the human oral microbiota.

### Fast-evolving microbes dominate the oral microbiome

One intriguing observation from the human oral microbiome is the high proportion of rapidly evolving species, which hinders the identification of species-specific marker genes for marker-based profiling. We were able to identify species-specific markers for fewer than half of the oral microbial species. Similarly, in a previous study of the gut microbiome, we identified 619 species with high divergence within the *Collinsella* genus, accounting for ∼11% of gut microbial species^7^.

To systematically identify rapidly evolving species in the oral microbiota, we analyzed the distribution of average phylogenetic distances to the 10 nearest neighbors for each species. Rapidly evolving species are expected to exhibit shorter phylogenetic distances to their close relatives compared to those with typical evolutionary rates. Consistently, we observed that species lacking species-specific marker genes had significantly lower distance scores, whereas species with such markers fell within a higher distance range (**Extended Data Fig. 3a**). This positive association between phylogenetic distance and the availability of marker genes highlights the limitations of marker-based profiling for rapidly evolving species. Using a phylogenetic distance threshold derived from the mean of the distance scores, which effectively separates the two modes, we identified 2,637 fast-evolving species, accounting for 77.0% of the 3,426 oral microbial species. This large proportion of fast-evolving species, which lack species-specific markers, underscores the unsuitability of marker-based profiling for the oral microbiome.

From the HROM phylogenetic tree, we identified six clades enriched in fast-evolving species: the phylum Patescibacteria, the family Campylobacteraceae, and four genera (*Rothia*, *Lancefieldelta*, *Oribacterium*, and *Streptococcus*) (**Extended Data Fig. 3b**). We hypothesized that an accelerated evolutionary rate is associated with increased genetic variation, anticipating a negative correlation between phylogenetic distance to close species and single nucleotide variant (SNV) density, consistent with findings from our previous study^7^. Among 514 species with more than 10 conspecific genomes, we observed this inverse correlation, indicating higher genetic variability in fast-evolving species (**Extended Data Fig. 3c**). Bacterial genome evolution is driven not only by genetic variability but also by adaptability to new environments. Accessory genes, which often provide selective advantages, may play a significant role in adaptive evolution^30^. Consistent with this, we observed a correlation between phylogenetic distance to close species and the proportion of core genes in the pangenome for the 514 species, suggesting larger accessory genomes in fast-evolving species (**Extended Data Fig. 3d**). Together, these findings confirm that species with shorter phylogenetic distances to close relatives exhibit faster evolutionary rates compared to others.

### HROM reveals a novel functional repertoire of the human oral microbiome

To explore the functional landscape of the human oral microbiome, we clustered species within each phylum based on Jaccard distances derived from presence/absence profiles of proteins, using representative genomes clustered at 50% sequence identity. This suggests that the phylogenetic relationships among HROM species align with their functional similarities (**Fig. 3a**). Notably, HROM cataloged many novel protein families that were not previously identified in bacterial protein databases.

**Fig. 3.**
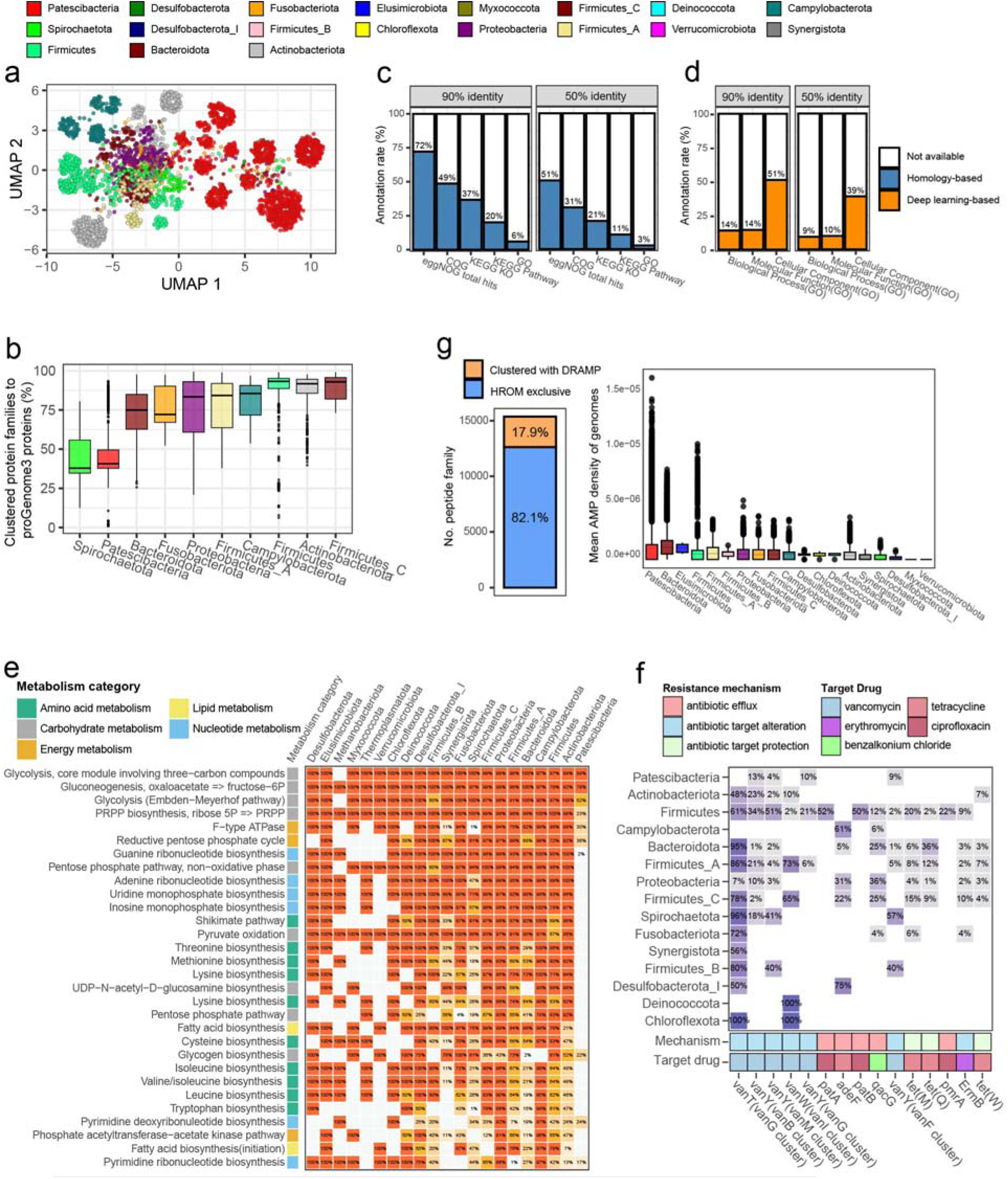
Functional and metabolic landscape of the HROM. **a**, UMAP visualization of 3,426 HROM species based on Jaccard distances derived from protein presence/absence profiles. Protein sequences from representative genomes were clustered into families at 50% sequence identity. **b**, The boxplot illustrates the percentage of proteins from representative genomes of each species clustered with proGenomes3 proteins at 50% identity, grouped by phylum. **c**, Annotation rates of HROM protein families at 90% and 50% sequence identity for Cluster of Orthologs Groups (COG), KEGG Ortholog (KO), KEGG pathway, and Gene Ontology (GO) terms using a homology-based method, eggNOG-mapper. **d**, Annotation rates of HROM protein families at 90% and 50% sequence identity for GO biological processes, molecular functions, and cellular components terms using a deep learning-based method, DeepGOPlus. **e**, Heatmap of the top 20 most abundant KEGG metabolic modules across HROM phyla, categorized into amino acid metabolism, carbohydrate metabolism, energy metabolism, lipid metabolism, and nucleotide metabolism. Numbers indicate the prevalence of each module within phyla. A metabolic module was considered present if it was identified in at least one conspecific genome within the same species. **f**, Heatmap of the top 15 most abundant antibiotic resistance genes with their mechanisms and target drugs across HROM phyla. The proportion of genomes encoding resistance mechanisms such as efflux, target alteration, and target protection is shown for each phylum and drug type. **g**, Summary statistics of antimicrobial peptides (AMPs) found in HROM. Left: Proportion of HROM AMPs clustered with those from Database of Antimicrobial Peptides (DRAMP) compared to HROM-exclusive AMPs. Right: Boxplot showing the mean AMP density of genomes across phyla, highlighting the prevalence of AMPs within each taxonomic group. (b, g) Boxplot elements: center line (median), box edges (25th and 75th percentiles), whiskers (1.5× interquartile range).

When comparing HROM proteins from representative species clustered at 50% sequence identity with those in proGenomes3^31^, the most comprehensive genome catalog containing nearly a million publicly available bacterial genomes, we identified a substantial number of unique proteins that did not cluster with any previously known bacterial proteins (**Fig. 3b**). This highlights the substantial contribution of HROM to expanding the repertoire of bacterial protein families, particularly for the phyla Spirochaetota and Patescibacteria.

We then identified protein-coding genes from 145,149 MQ genomes using Prokka^32^. The predicted protein sequences were clustered at 100%, 95%, 90%, 70%, and 50% sequence identity using MMseqs2^33^, yielding approximately 76.2, 15.2, 8.5, 2.7, and 2.0 million protein clusters, respectively. We performed functional annotation for these protein families using the homology-based method eggNOG-mapper^34^, which assigns various functional terms including Cluster of Ortholog Groups (COG)^35^, KEGG Orthology (KO)^36^, KEGG Pathway^36^, Gene Ontology (GO)^35^. We observed that most protein families lacked functional annotations across these terms (**Fig. 3c**). Notably, GO annotations covered only 6% and 3% of protein families at 90% and 50% sequence identity threshold, respectively. To address this limitation, we applied DeepGOPlus^37^, a deep learning-based GO term prediction tool, which substantially improved the annotation rate, particularly for cellular component terms (**Fig. 3d**).

The landscape of KEGG pathways revealed that glycolysis and gluconeogenesis were highly prevalent across all HROM phyla, with other metabolic pathways also common in most phyla, while phylum Patescibacteria exhibited a marked depletion of metabolic modules (**Fig. 3e**, **Supplementary Table 7**). In addition, amino acid and nucleotide metabolism were notably absent in phyla Myxococcota and Verrucomicrobiota, and energy metabolism was lacking in phyla Methanobacteriota and Thermoplasmatota. The landscape of antibiotic resistance genes showed that glycopeptide resistance gene targeting vancomycin (vanT, vanY and vanW) were prevalent in oral microbiome (**Fig. 3f**). Diverse defense mechanisms against phage infection were found across phyla, with restriction modification systems were most prevalent **(Extended Data Fig. 4a**). The landscape of biosynthetic gene clusters (BGC) showed that categories of secondary metabolites RiPP-like, arylpolyene, T3PKS, NRPS were relatively more prevalent in HROM **(Extended Data Fig. 4b)**.

Many small peptides encoded by bacterial small open reading frames (sORFs) also exhibit broad-spectrum antimicrobial activates and play key roles in microbial interactions^38^. Using Macrel^39^, we predicted anti-microbial peptides (AMPs) from HROM, identifying 15,368 AMPs, of which only 2,744 (∼17.9%) clustered at family level with 26,108 peptides in the DRAMP^18^, a public repository of AMPs **(Fig. 3g, Supplementary Table 8)**. This highlights that HROM includes a substantial number of previously unreported AMPs. The density of AMP per genome was relatively low overall, with phyla Patescibacteria and Bacteroidota showing comparatively higher mean densities.

In conclusion, HROM unveils an extensive repertoire of functions in the human oral microbiome by identifying novel protein families and antimicrobial peptides. With improved functional annotation and comprehensive profiling of the antibiotic resistome, phage defensome, and secondary metabolism gene clusters, HROM serves as a valuable resource for facilitating functional studies of the oral microbiome.

### Distinct genomic and functional features of oral Patescibacteria unveiled by HROM

Intriguingly, we collected 7,321 NC genomes and 1,402 species from the phylum Patescibacteria, a member of the Candidate Phyla Radiation (CPR) group within the human oral microbiome. Of these, 7,107 were assembled using our HROM pipeline, and 2,903 genomes correspond to 1,137 (81%) novel species unclassified at the species level by GTDB taxonomy. Previous studies using concatenated protein markers have identified the CPR as a monophyletic group encompassing numerous phyla^40,41^. However, the GTDB taxonomy, a standardized bacterial classification system based on genome phylogeny and rank normalization, has reclassified CPR as a single phylum, Patescibacteria^42^. Patescibacteria are characterized by ultrasmall cell sizes, reduced genomic capabilities, small genome sizes, limited metabolic capacity^20^, and an epi-symbiotic lifestyle^43^ with host bacteria. Their obligate symbiosis is thought to drive rapid evolutionary processes, contributing to their high genetic divergence^44^ (**Extended Data Fig. 3b**).

Notably, oral Patescibacteria species exhibit smaller genome sizes compared to other oral bacterial species (**Extended Data Fig. 5a**). Representative genomes of Patescibacteria lack 21 of the 120 universal single-copy genes used as GTDB markers, with these 21 markers showing a maximum prevalence of only 1.5% (**Extended Data Fig. 5b, Supplementary Table 9**). In contrast, the mean prevalence of all GTDB markers across other oral bacterial species is 93.19%, with a minimum of 65.17%, highlighting the significantly reduced genomic functionality of oral Patescibacteria. The division and cell wall (*dcw*) cluster, comprising 17 genes involved in peptidoglycan precursor synthesis, is reported to be highly conserved across the bacterial domain, including the CPR superphylum^45^. Interestingly, oral CPR bacteria have retained most of the genes in the *dcw* cluster (**Extended Data Fig. 5c**), suggesting that the reduced genome size reflects an inherent property of oral Patescibacteria rather than an artifact of incomplete genome assembly.

Previously, this group lacked isolated representative genomes until recent studies^46^. The protein content of CPR bacterial genomes is distinct from that of other bacterial genomes^47^, highlighting their functional uniqueness. The functional distinctiveness of oral Patescibacteria was demonstrated by their separate clustering from other oral bacterial species based on protein family profile distances (see **Fig. 3a**). Consistent with their reduced genome sizes, Patescibacteria are comparably depleted in metabolic capabilities relative to other oral bacterial species (**Fig. 4a**). However, oral Patescibacteria genomes exhibit a higher proportion of genes encoding transmembrane proteins compared to other species (**Fig. 4b, Extended Data Fig. 5d**). This suggests that oral Patescibacteria genomes may prioritize genes related to transport, consistent with their parasitic lifestyle. Using KO and COG terms, we identified several genes in oral Patescibacteria species that are enriched for biofilm formation functions compared to other species: VpsQ (K20950) is related to the production of exopolysaccharide, which are crucial for mature biofilm formation^48^; resuscitation-promoting factor (K21688) is crucial for biofilm formation in some species^49^ or containing catalytic domains that mediates biofilm formation^50^; type IV pilus-related genes such as pilM (K02662, COG4972) and VirB6 (COG3704) indirectly contribute to biofilm formation^51–53^ (**Extended Data Fig. 5e-f Supplementary Table 10**). These findings suggest that oral Patescibacteria species play a significant role in biofilm formation within the oral microbiota, emphasizing their importance in community structure and function.

**Fig. 4.**
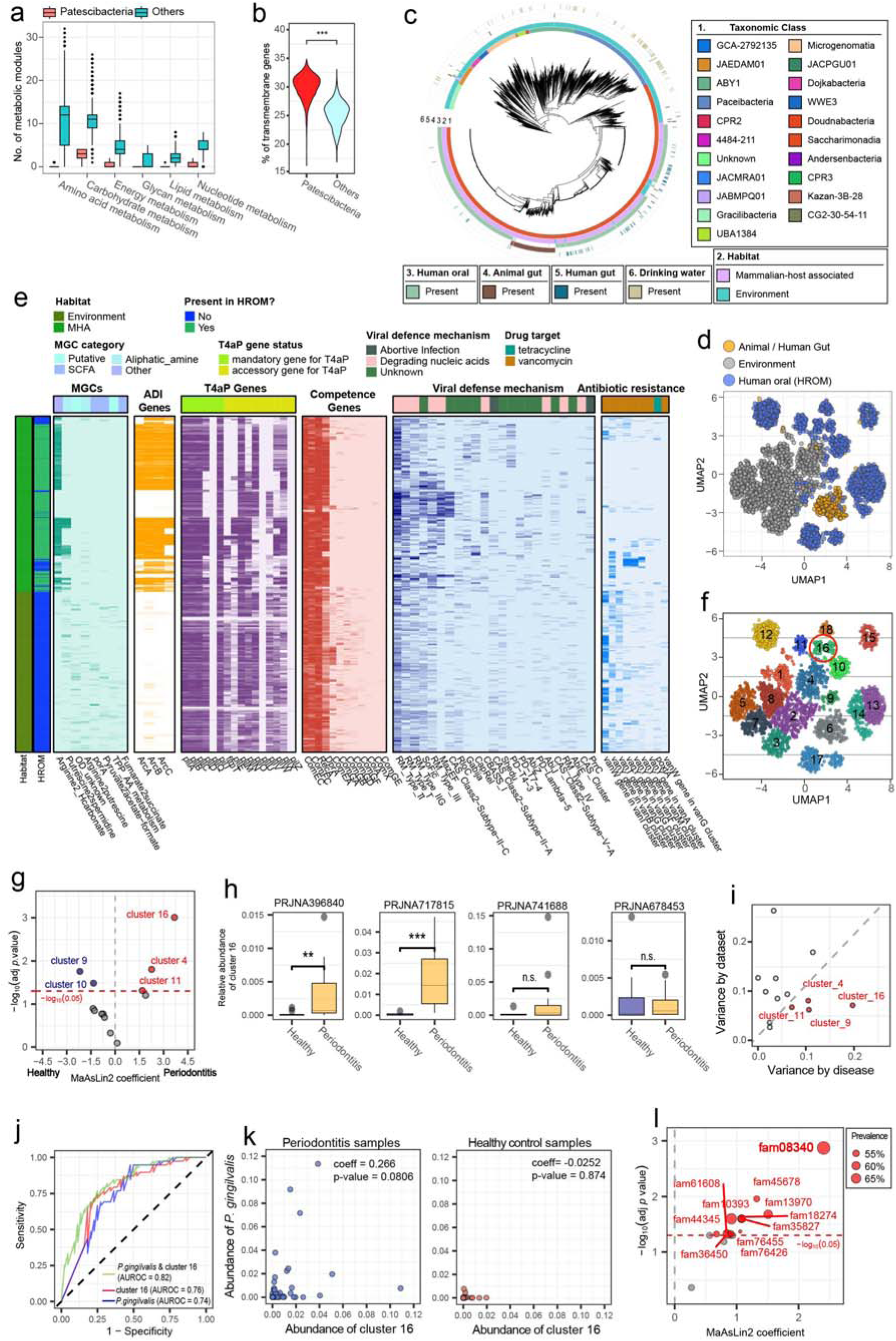
Functional, phylogenetic, and disease-associated analysis of oral Patescibacteria. **a**, Comparison of the number of metabolic modules in Patescibacteria versus other phyla, categorized into amino acid metabolism, carbohydrate metabolism, energy metabolism, glycan metabolism, lipid metabolism, and nucleotide metabolism. **b**, Violin plot showing the percentage of transmembrane genes in Patescibacteria compared to other phyla. Statistical significance was assessed using a two-tailed Mann-Whitney U test (*** *P* < 0.001). **c**, Phylogenetic tree of 2,963 Patescibacteria species, with outer tracks indicating taxonomic class, habitat, and presence in various environments, including human oral cavity, gut, animal gut, and drinking water. **d**, UMAP visualization of protein family profiles across 2,963 Patescibacteria species representative genomes based on the Jaccard distance, showing clustering by environmental sources (e.g., human oral cavity and gut, animal gut, environment). **e**, Heatmap showing the distribution of metabolic gene clusters (MGCs), ADI genes, T4aP genes, competence genes, viral defense mechanisms, and antibiotic resistance genes across 2,963 Patescibacteria species. Colors represent habitat types and gene presence. **f**, UMAP visualization of Patescibacteria species clusters based on protein family profiles similarity. Each cluster is numbered and color-coded, with cluster 16 highlighted by a red circle. **g**, Identification of differentially enriched Patescibacteria clusters in periodontitis samples using MaAsLin2. Clusters meeting the adjusted p-value threshold (*P* ≤ 0.05, Bonferroni-Hochberg correction) are highlighted in red, with the horizontal red line indicating the significance threshold. Cluster 16 is strongly associated with periodontitis. **h**, Relative abundance of cluster 16 across four independent periodontitis datasets, showing significant enrichment in two out of four periodontitis samples compared to healthy controls. Statistical significance was assessed using a two-tailed Mann-Whitney U test (*** *P* < 0.001, ** *P* < 0.01, n.s. = not significant). **i**, Variance explained by dataset (y-axis) versus disease label (x-axis) for different clusters of Patescibacteria species in periodontitis-associated datasets, with clusters showing greater disease-associated variance highlighted in red. **j**, Receiver operating characteristic (ROC) curve for periodontitis classification based on the batch-corrected relative abundance of *P. gingivalis* (blue), cluster 16 (red), and their combined abundance (green). The area under the ROC curve (AUROC) scores are indicated. **k**, Comparison of the batch-corrected relative abundance of cluster 16 and *P. gingivalis* in healthy and periodontitis samples. The Pearson correlation coefficient and *p*-value are indicated. **l**, Enriched protein families in cluster 16 associated with periodontitis. The size of each dot represents prevalence, and significant families are labeled with MaAsLin2 coefficients and adjusted *p*-values threshold (*P* ≤ 0.05, Bonferroni-Hochberg correction) are highlighted in red, with the horizontal red line indicating the significance threshold. (a, h) Boxplot elements: center line (median), box edges (25th and 75th percentiles), whiskers (1.5× interquartile range).

Given our extensive collection of CPR group bacterial genomes from the human oral microbiome, we investigated the phylogenetic and functional divergence of mammalian host-associated (MHA) CPR bacteria, including oral Patescibacteria, compared to those inhabiting environmental niches such as soil and water. To achieve this, we gathered CPR bacterial genomes from other isolation sources to analyze the functional and phylogenetic traits of oral Patescibacteria in the context of other known Patescibacteria (**Supplementary Table 11**). Genomes belonging to the phylum Patescibacteria were retrieved from previous studies^54^ and public repositories and were filtered using criteria identical to those applied to the HROM. This effort resulted in the collection of 3,721 genomes, which were subsequently de-replicated into 2,963 species clusters (**Fig. 4c**). Notably, 84.6% (2,507 species) were singletons, likely reflecting the rapid speciation characteristic of this group. Among the species-level genomes, 1,423 (48%) originated from environmental samples, whereas 1,540 (52%) were derived from host-associated sources. Phylogenetic analysis revealed that Patescibacteria species from the human oral cavity are phylogenetically closer to genomes isolated from the human gut, skin, and other mammalian hosts compared to those isolated from environmental sources such as soil and water. Interestingly, over 98% of the 1,553 MHA Patescibacteria species were classified within the class Saccharimonadia, highlighting their taxonomic distinctness and adaptation to mammalian host environments. Using a dataset of 22,977 protein families from the 2,963 representative genomes of Patescibacteria^47^, we performed clustering based on the Jaccard distance between protein family profiles. This analysis showed clear, non-overlapping clusters among species associated with different environments (**Fig. 4d**), further supporting the functional adaptation of Patescibacteria genomes to their respective environments.

We examined functions that were more prevalent in MHA or environmental Patescibacteria (**Supplementary Table 12, Extended Data Fig. 5g**). Among the identified genes, the ribonucleoside-triphosphate reductase gene (*nrdD*), which has previously shown high phylogenetic diversity in MHA Patescibacteria^55^, was found to be more prevalent in MHA genomes. This gene enables survival in anaerobic conditions, such as those present in the oral cavity or gut. Similarly, L-lactate dehydrogenase (*ldh*), another gene previously reported to be unique to the MHA group^55^, was more prevalent in MHA Patescibacteria, particularly in oral species. The *ldh* gene is a signature of oral bacteria, facilitating the production of lactate, a highly abundant metabolite in dental plaque. Additionally, ABC-type transport systems were enriched in MHA Patescibacteria, indicating their enhanced ability to import and export essential resources necessary for survival in host environments (**Extended Data Fig. 5h**).

We further examined pathways relevant to CPR bacteria. The arginine deiminase (ADI) system, known to protect both CPR bacteria and their host bacteria from acid stress^56^ and essential for the growth of CPR bacteria^57^, was found to have core genes (*ArcA, ArcB, ArcC*) and the associated metabolic gene cluster (MGC) highly prevalent in MHA Patescibacteria (**Fig. 4e**). In contrast, the Type IVa pilus system (T4aP), which is a known trigger for epi-symbiotic relationships with host bacteria^58^, was prevalent across all Patescibacteria. Similarly, competence genes such as *comEC* and *comFC*, required for the processing of extracellular DNA and facilitating potential horizontal gene transfer (HGT)^59^, were found across all Patescibacteria. Lastly, MHA Patescibacteria demonstrated a comparatively more diverse array of viral defense systems but exhibited a depletion of antibiotic resistance genes relative to their environmental counterparts. Notably, oral CPR bacteria lacking the ADI system exhibited a higher abundance of phage defense systems, suggesting distinct subgroups of oral CPR bacteria that enhance host bacteria ability to either resist acid stress or defend against phage infection.

### Unveiling an oral CPR subclade associated with periodontitis

CPR bacteria has been implicated for periodontitis, a serious gum disease characterized by inflammation and eventual destruction of the bone supporting the teeth^44^. However, no specific CPR subclade or functional elements directly associated with periodontitis have been identified. Leveraging our extensive collection of oral CPR species genomes, we aimed to identify CPR bacterial species linked to periodontitis.

Given the high similarity in gene contents among CPR species due to rapid speciation, univariate analysis of individual species may be suboptimal. To address this, we clustered 18 groups of oral CPR bacterial species based on Jaccard distances of protein family presence profiles (**Extended Data Fig. 6a**) and tested their association with periodontitis using publicly available WMS data. This analysis identified cluster 16 CPR bacteria (**Fig. 4f, Supplementary Table 13**) as the most enriched group in saliva-sampled periodontitis cohorts (**Fig. 4g**). The relative abundance of cluster 16 CPR bacteria was significantly higher in two of four periodontitis cohorts and showed a similar but non-significant trend in another cohort (**Fig. 4h**). Despite the limited number of datasets, disease states explained a higher proportion of variance than dataset-specific factors (**Fig. 4i**). These findings suggest an association between the abundance of oral CPR bacteria in cluster 16 and periodontitis. Further, phylogenetic analysis revealed that cluster 16 CPR bacteria comprise a subclade of the genus *Nanosynbacter* (**Extended Data Fig. 6b-c**).

A well-known oral bacterial species implicated in periodontitis is *Porphyromonas gingivalis*. We observed that the relative abundance of *P. gingivalis* within the oral microbiome predicts the occurrence of periodontitis with an area under the receiver operating characteristic curve (AUROC) of 0.74 (**Fig. 4j**). Notably, the relative abundance of cluster 16 CPR bacteria provided slightly better predictive performance (AUROC = 0.76) than *P. gingivalis*. Moreover, the combined relative abundance of *P. gingivalis* and cluster 16 CPR bacteria further improved periodontitis prediction, achieving an AUROC of 0.82. This finding suggests that *P. gingivalis* and cluster 16 CPR bacteria are independently associated with periodontitis. Supporting this, we observed a relatively low correlation between their relative abundances in periodontitis patients (**Fig. 4k**). These oral microbiomes of patients were generally characterized by either abundant *P. gingivalis* or abundant cluster 16 CPR bacteria.

We found fam08340 as the most prevalent protein family among cluster 16 species **(Fig. 4l, Extended Data Fig. 6d-e**). This protein family lacks functional annotation, and no homologous proteins with known functions were identified. Analysis of the ∼150 amino acid N-terminal region of fam08340 revealed a signal peptide linked to transmembrane region. To explore potential structure-based functions, we predicted the 3-dimensional structure of the outer-membrane region of the protein using ESMFold^60^ and conducted a structural homology search against AlphaFold database(UniProt50)^61^ with Foldseek^62^ (**Extended Data Fig. 6f**). Among the top hits, we identified polycystic kidney disease (PKD) domain-containing proteins (**Extended Data Fig. 6g**). The PKD domain, an 80-90 amino acid module characterized by a β-sandwich fold, facilitates ligand-binding interactions and is critical in the pathogenesis of autosomal dominant polycystic kidney disease (ADPKD)^63^. Notably, a carboxypeptidase from *P. gingivalis* contains three tandem PKD domains, which are implicated in its role in disease pathogenesis^64^. This suggests that fam08340 of cluster 16 CPR bacteria may similarly contribute to periodontitis through mechanisms analogous to the carboxypeptidase of *P. gingivalis*.

In summary, leveraging a substantially expanded catalog of CPR species genomes and their proteins, we identified a subclade of oral CPR bacteria implicated in periodontitis and uncovered proteins potentially involved in its pathogenesis.

### Functional divergence and taxonomic overlap between oral and gut microbiomes

The human oral cavity and gut are physiologically distinct yet directly interconnected body sites. This interconnection may lead to the divergence of microbiomes between two sites while allowing microbial exchange and cross-site influence^65^. For example, a significant number of oral bacteria are transferred to the gut daily, primarily through the ingestion of saliva^66–68^. To explore the taxonomic and functional relationship between the microbiomes of the oral cavity and gastrointestinal tract, we compared HROM with HRGM^7^, a human reference gut microbial genome catalog. To improve reliability of this study, we utilized HRGM2, an improved version of HRGM exclusively composed of high-quality genomes from 4,824 species (manuscript submitted).

We first observed a substantially smaller genome size in the oral microbiota compared to the gut microbiota (**Fig. 5a**). Since both HROM and HRGM2 contain only high-quality genomes with completeness greater than 90%, this difference cannot be attributed to the quality of the genome catalogs. This skewed distribution toward smaller genome sizes appears to be largely attributable to oral CPR bacteria.

**Fig. 5.**
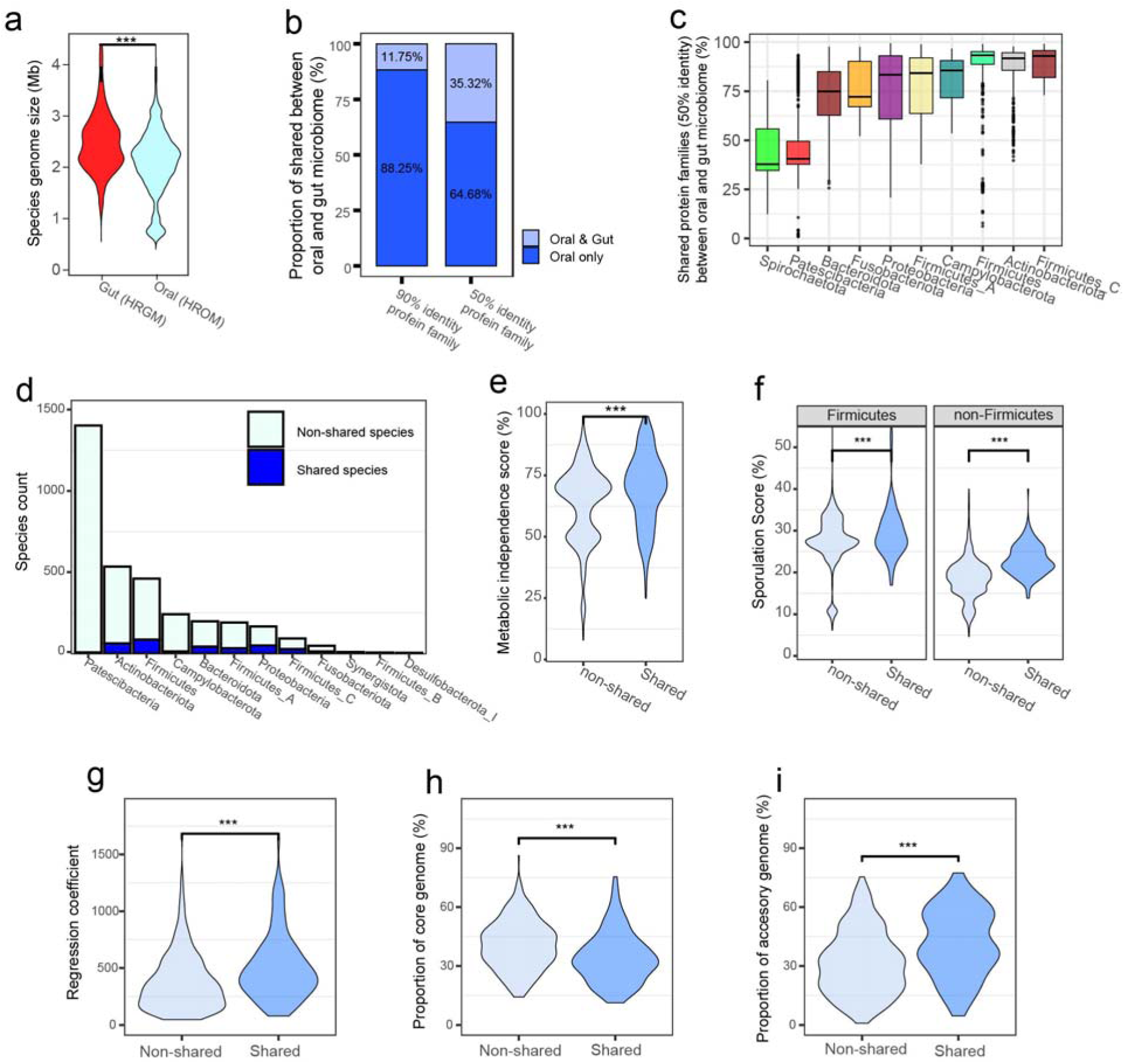
Functional analysis of oral-gut shared bacterial species. **a**, Violin plot comparing the genome size of species in the gut (HRGM2) versus the oral cavity (HROM). Statistical significance was assessed using a two-tailed Mann-Whitney U test (*** *P* < 0.001). **b**, Proportion of shared protein families between oral and gut microbiomes at 90% and 50% sequence identity, showing the percentages of protein families exclusive to the oral microbiome and those shared with the gut microbiome. **c**, Boxplot of the proportion of shared protein families (50% sequence identity) between oral and gut microbiomes across major phyla. Boxplot elements: center line (median), box edges (25th and 75th percentiles), whiskers (1.5× interquartile range). **d**, Species count for oral-gut shared and non-shared species across major phyla. Shared species are defined as those found in both microbiomes. **e**, Violin plot comparing the metabolic independence score (%) between oral-gut shared and non-shared species. Patescibacteria were excluded due to their low metabolic independence and minimal oral-gut sharing to prevent bias. Statistical significance was assessed using a two-tailed Mann-Whitney U test (*** *P* < 0.001). **f**, Violin plot comparing sporulation scores (%) between oral-gut shared and non-shared species, grouped by Firmicutes and non-Firmicutes, with each species’ score determined by the maximum value among its conspecific genomes. Statistical significance was assessed using a two-tailed Mann-Whitney U test (*** *P* < 0.001). **g**-**i**, Violin plots comparing the regression coefficient of pangenome openness, with higher coefficients indicating a more open pangenome (**g**), the proportion of core genome (%) (**h**), and the proportion of accessory genome (%) (**i**) between oral-gut shared and non-shared species (based on 125 shared and 389 non-shared species). Statistical significance for these panels was assessed using a two-tailed Mann-Whitney U test (*** *P* < 0.001).

Next, we compared protein families at 90% and 50% sequence identity and found that the majority (∼88% and ∼65%, respectively) were unique to the oral microbiome (**Fig. 5b**), indicating that the oral microbiome is functionally distinct from the gut microbiome. The overlap of protein families between the two microbiomes varied among phyla; notably, Spirochaetota and Patescibacteria exhibited particularly low overlap, which could suggest that these two phyla might represent the most functionally divergent taxonomic groups in the oral microbiome. (**Fig. 5c**). These findings highlight that the symbiotic microbes of the oral cavity and gut have adapted to their distinct environments through functional divergence.

The anatomical connection between the oral cavity and gastrointestinal tract suggests the potential for shared microbial species. Based on a criterion of 95% ANI (average nucleotide identity) of genomes, we identified 330 species present in both the oral and gut microbiomes (**Supplementary Table 14**). Most of these oral-gut shared species were classified within the phylum Firmicutes, followed by Actinobacteriota and Proteobacteria (**Fig. 5d**).

We hypothesized that these shared species might possess functional characteristics that enable their ubiquitous presence. Specifically, we propose that microbes capable of inhabiting multiple environments are more likely to be metabolically independent. To test this, we used a metabolic independence score, calculated as the summation of the completeness of 33 KEGG metabolic modules known to be enriched in bacteria identified as good colonizers during fecal microbiota transplantation (FMT)^69^. Comparing the metabolic independence scores of the oral-gut shared species with those of non-shared species revealed that shared species exhibited significantly higher metabolic independence (**Fig. 5e**).

Additionally, shared species are likely to survive in the harsh gastric environment during transmission from the oral cavity to the gut. Since endospore formation enhances bacterial survival in adverse conditions, we evaluated the sporulation capacity of these species using a previously established sporulation score, which is based on the presence of 66 sporulation-predictive genes in bacterial genomes^70^. Shared species showed significantly higher sporulation scores than non-shared species (**Fig. 5f**), supporting their enhanced probability of successful transmission from the oral cavity to the gut.

Furthermore, we found that oral-gut shared species exhibit a higher tendency for an open pangenome compared to non-shared species. This was evidenced by a higher pangenome regression coefficient (**Fig. 5g**), a lower proportion of core genome (**Fig. 5h**), and a higher proportion of accessory genome (**Fig. 5i**). These findings align with previous observations that bacteria with open pangenomes are more adaptable to diverse environments.

### Ectopic oral species abundance in gut microbiota linked to systemic diseases

Recent evidence suggests that oral microbes influence not only oral diseases but also a range of systemic disorders, including colorectal cancer (CRC), inflammatory bowel disease (IBD), and cardiovascular diseases^3^. Although the roles of oral microbes in systemic diseases are not yet fully understood, growing evidence indicates that oral bacteria can migrate into the gastrointestinal tract and contribute to pathological processes there^65^. Based on this, we hypothesized that some of the 330 oral-gut shared species may be ectopic oral bacteria—species typically residing in the oral cavity but occasionally detected in non-typical sites such as the gut—rather than dual-site residents. To distinguish ectopic oral species from oral-gut shared species, we focused on those prevalent in healthy oral microbiomes but not in healthy gut microbiomes. Specifically, we selected shared species present in more than 50% of healthy oral microbiome samples but less than 25% of healthy gut microbiome samples on average across cohorts (**Fig. 6a**, **Supplementary Tables 15–16**). This approach identified 42 ectopic oral species (**Supplementary Table 17**).

**Fig. 6.**
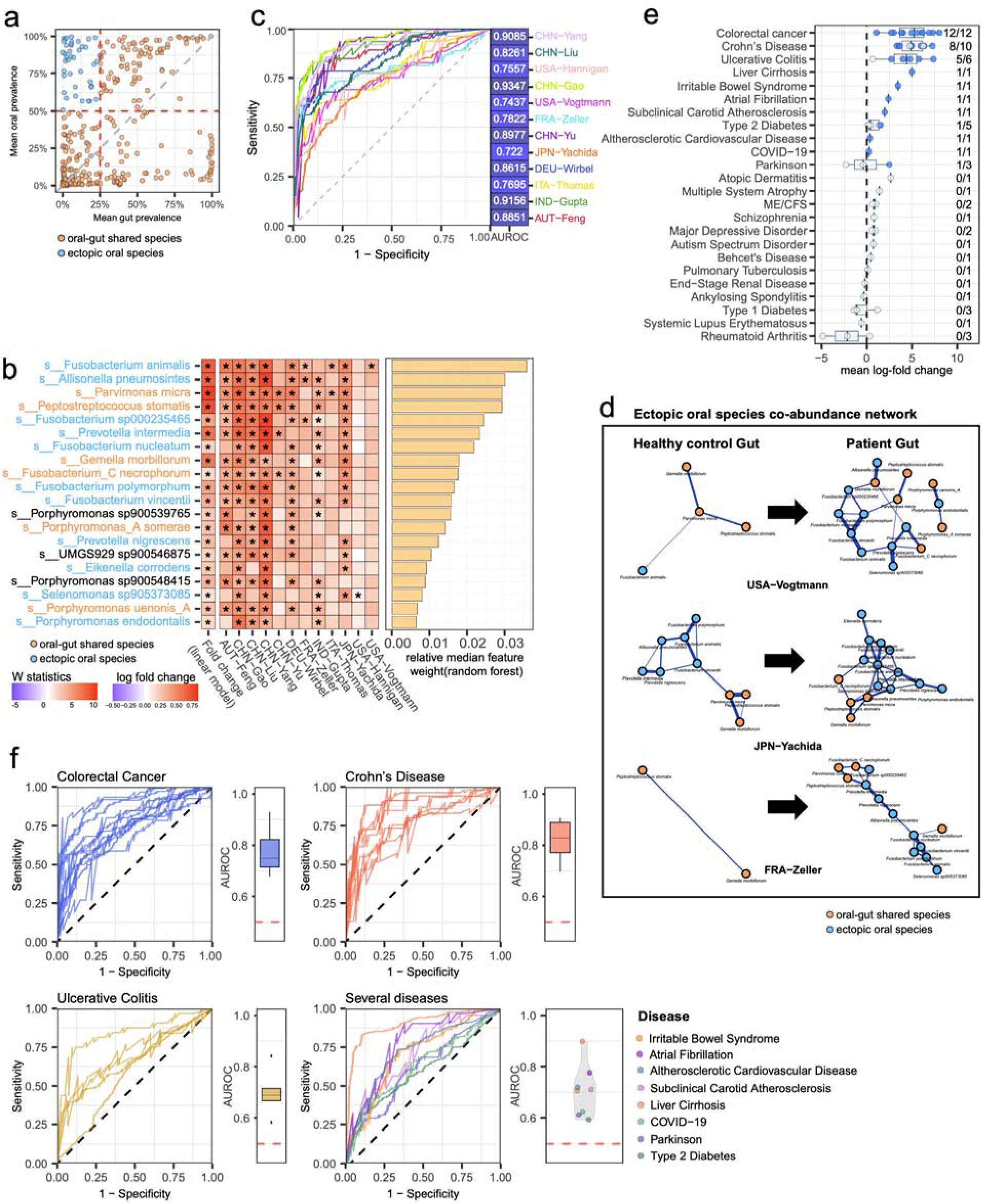
Disease association of ectopic oral species. **a**, Scatterplot showing the mean prevalence of 330 oral-gut shared species (orange) across 9 healthy gut samples (x-axis) and 14 healthy oral samples (y-axis). Forty-two ectopic oral species (blue) are defined as those present in more than 50% of healthy oral samples but less than 25% of healthy gut samples. The prevalence thresholds are indicated by red lines. **b**, Heatmap displaying the top 20 bacterial species identified by random forest models as important features for colorectal cancer (CRC) classification. The left heatmap shows log-fold changes derived from linear models using SIAMCAT, with statistical significances determined by FDR-adjusted p-values (* adjusted *P* ≤ 0.05). The central heatmap presents W statistics from ANCOM-BC analysis across 12 CRC cohorts, with stars denoting statistically significant associations (* adjusted *P* ≤ 0.05) based on Holm-Bonferroni-corrected p-value. The bar chart (right) displays the relative median feature weight of each species extracted from random forest model. **c**, Receiver operating characteristic (ROC) curves and area under the ROC (AUROC) values for CRC prediction across 12 cohorts based on the sum of relative abundances of the 17 oral species among the top 20 key species for CRC classification. **d**, Co-abundance network of 17 oral species among the top 20 key species for CRC classification in healthy (left) and patient (right) gut microbiome samples across different cohorts (USA-Vogtmann, JPN-Yachida, FRA-Zeller). Orange and blue nodes represent oral-gut shared and ectopic oral species, respectively. Thicker edges in patient networks indicate stronger co-abundance relationships. **e**, Mean log-fold change of sum of relative abundance of 42 ectopic oral species in gut microbiomes associated with various systemic diseases, including colorectal cancer, Crohn’s disease, ulcerative colitis, and cardiovascular diseases. Disease association is indicated by the fraction of cohorts showing a significant increase in abundance (two-tailed Mann-Whitney U test, *P* ≤ 0.05). Numbers on the right side indicate the number of datasets with significant enrichment. ME/CFS: Myalgic Encephalomyelitis/Chronic Fatigue Syndrome. **f**, ROC curves and AUROC values for disease classification using the sum of relative abundance of 42 ectopic oral species for colorectal cancer, Crohn’s disease, ulcerative colitis, and several other diseases. Boxplots display AUROC values across cohorts, with significant predictive performance for intestinal diseases and moderate performance for other diseases. (e-f) Boxplot elements: center line (median), box edges (25th and 75th percentiles), whiskers (1.5× interquartile range).

To validate these ectopic oral species, we focused on the gut microbiome of CRC, an intestinal disease known to be influenced by oral bacteria such as *Fusobacterium nucleatum*^71^. We analyzed 12 publicly available CRC gut microbiome datasets (comprising 910 healthy individuals and 877 patients) taxonomically profiled with HRGM2 (**Supplementary Table 18-19**). Using microbial species abundance profiles from nine datasets for training and three datasets for validation, we constructed disease classifier with a random forest algorithm. Species were then prioritized for feature importance based on their random forest feature weights. Remarkably, 55% of the top 20 important species associated with CRC were identified as ectopic oral species (**Fig. 6b**). Furthermore, CRC prediction using the sum of the relative abundances of 17 oral species (oral-gut shared species and ectopic oral species) among the top 20 important species achieved a significant performance, with a mean AUROC value of 0.834 (**Fig. 6c**). Additionally, we observed that the co-abundance networks of these 17 oral species were enhanced in the gut microbiomes of CRC patients compared to healthy controls (**Fig. 6d**, **Extended Data Fig. 7**), potentially indicating the co-translocation of these oral microbial species into the gut microbiome is associated with CRC.

Next, we tested whether the relative abundance of these 42 ectopic oral species increased in the gut microbiome of systemic diseases, including CRC. To do this, we measured the fold change in the sum of relative abundance of the ectopic oral species in patient fecal microbiomes compared to healthy controls across multiple disease-associated cohorts, encompassing 24 disease types (**Supplementary Table 18-19**). We observed a substantial and consistent increase in the relative abundance of ectopic oral species in gut microbiome samples from patients with intestinal diseases (e.g., CRC, Crohn’s disease, ulcerative colitis, irritable bowel syndrome), cardiovascular diseases (e.g., atrial fibrillation, subclinical carotid atherosclerosis, atherosclerotic cardiovascular disease), and liver cirrhosis (**Fig. 6e**). Furthermore, the sum of the relative abundance of the 42 ectopic oral species in the gut microbiome showed high predictive power for these diseases (**Fig. 6f**). For other diseases such as COVID-19, Parkinson’s disease, and type 2 diabetes, we also observed an increased sum of relative abundance of ectopic oral species in patient gut microbiomes compared to healthy controls. However, the effect sizes were smaller or inconsistent across cohorts, resulting in only moderate disease prediction accuracy.

In summary, we identified 42 ectopic oral species by comparing oral and gut bacterial reference genomes and analyzing their differential prevalence between healthy oral and gut microbiome samples. We uncovered their significant associations with various systemic diseases, particularly intestinal diseases, cardiovascular diseases, and liver diseases. These findings highlight the value of the genomic catalog of the oral microbiome in advancing our understanding of how oral microbes influence systemic health.

## Discussion

In this study, we present the most comprehensive genomic catalog of the human oral microbiome to date, HROM, including 72,641 NC genomes assembled from 3,426 species, of which 2,019 (59%) are novel. This represents a substantial expansion over preexisting catalogs, increasing the classification rate of metagenomic sequence reads significantly. This improvement facilitates more accurate taxonomic profiling of WMS data from oral microbiota samples. Notably, we identified a high prevalence of fast-evolving species among oral microbiota, which limits the identification of reliable species-specific markers. This challenge results in suboptimal accuracy for marker-based profiling methods like MetaPhlAn4. Therefore, we recommend DNA-based methods, such as Kraken-Bracken, for species profiling of oral microbiota. To ensure versatility, we have provided custom genome databases for both DNA-based and marker-based profiling methods in the accompanying companion webserver.

Unexpectedly, we identified 1,137 novel CPR bacterial species from oral microbiota through metagenome assembly, establishing Patescibacteria as one of the most prevalent phyla in the human oral microbiota. Functional and phylogenetic analyses revealed that oral Patescibacteria form a clade distinct from environmental Patescibacteria. This suggests that CPR bacteria, initially introduced from exposed environments like groundwater (a major habitat for CPR bacteria), have adapted to the host environment through functional repurposing of their genomes. Functional analysis of oral Patescibacteria genomes revealed key roles, including (i) biofilm formation, aiding microbial communities in the oral cavity; (ii) protection against acid stress, supporting host bacteria survival; (iii) nutrient supply for host bacteria through specialized transport systems; (iv) defense against phage infections, through a diverse and abundant defensome. Given their epi-symbiotic relationships with host bacteria, the diversification of oral CPR bacterial species may be driven by the diversity of their host bacteria. Identifying CPR-host species pairs and understanding their co-evolution represents an important avenue for future oral microbiota research. Additionally, we identified a novel oral CPR subclade associated with periodontitis and demonstrated that they complement *P. gingivalis* in predicting the disease. These findings expand our genomic understanding of oral pathogens and suggest that the oral CPR subclade may facilitate early diagnosis of periodontitis in clinical settings.

Although HROM database includes MQ genomes, we recommend using only NC genomes and species containing at least one NC genome (referred to as “NC genome species”). MQ genomes can lose up to 50% of their genomic content, which may result in inaccuracies in taxonomic and functional analyses. For instance, including MQ genomes may identify 1,687 additional species; however, 81.4% of these are singletons, likely representing technical artifacts rather than true biological diversity. We believe the exclusive use of high-quality genomes enhances the credibility of conclusions regarding functional divergence and taxonomic overlap between oral and gut microbiomes.

The transmission of oral bacteria into the gastrointestinal tract and their influence on systemic health has been suggested in previous studies. However, to our knowledge, no prior studies have systematically investigated oral bacteria associated with systemic diseases. By comparing HROM with HRGM, we first identified 330 bacterial species shared between the oral and gut environments. Among these, some may represent different subspecies adapted to each body site, while others likely consist of oral species found in atypical locations like the gut. These ectopic oral species may contribute to or exacerbate pathogenic conditions, such as inflammation. For the first time, we identified 42 ectopic oral species based on their prevalence in oral and gut samples from healthy individuals. We further demonstrated that their relative abundance in gut microbiota is predictive of intestinal diseases, cardiovascular diseases, and liver diseases. This pathogenic effect is likely mediated by systemic inflammation. In addition, we observed weak or inconsistent associations between the 42 ectopic oral species and diseases like COVID-19, Parkinson’s disease, and type 2 diabetes, which are also influenced by systemic inflammation. Consistent with this, a recent study of patient cohorts undergoing allogeneic hematopoietic cell transplantation or suffering from IBD revealed that the relative abundance of oral bacterial species in the gut increases as a marker of depleted gut bacterial diversity, while their absolute abundance does not significantly change^72^. Future research should confirm whether the 42 ectopic oral species similarly increase their relative abundance without a corresponding increase in absolute abundance. This distinction could provide deeper insights into their role in systemic health and disease.

This study has certain limitations. While we compiled the most comprehensive collection of WMS data from public repositories, the majority of the data originated from the USA and China. This geographic bias may impact the representativeness of the oral reference microbiome, particularly concerning ethnicity, geography, and lifestyle. Future updates should prioritize sampling from underrepresented regions to reduce these biases. Additionally, since most HROM genomes were assembled from short-read sequencing data, approximately 37% of species (1,265 out of 3,426) lack the 16S rRNA region, one of the most challenging regions to assemble. Future efforts employing long-read sequencing technologies could help resolve this limitation, enabling more complete genomic assemblies of the human oral microbiome. Moreover, the current version of HROM primarily focuses on bacterial genomes; however, future expansions will include fungal and viral genomes.

## Methods

### Collection of whole metagenome sequencing (WMS) data and bacterial isolate genomes from the human oral cavity

The WMS data were downloaded from public repositories, including ENA^12^, SRA^13^, and CNGBdb^14^. Each dataset underwent manual inspection to identify samples originating from the human oral cavity. Metadata, such as NCBI SRA biosample annotations, bioproject metadata, and its corresponding literature, were scrutinized to confirm sample origins. To collect pre-assembled genomes, we surveyed eHOMD (v3)^10^, GenBank^15^, RefSeq^16^, and PATRIC^17^. For eHOMD, genomes annotated as originating from non-oral environments were excluded. Biosample data, GCF/GCA accessions, and relevant literature were reviewed to verify that genomes were isolated from the human oral cavity. A similar process was applied to genomes from GenBank, RefSeq, and PATRIC, ensuring only genomes with verified isolate records in biosamples or literature were included. Pre-assembled MAGs derived from short-read sequencing were excluded, as most samples previously used for MAG assembly were reanalyzed in this study. Additionally, representative genomes from the oral bacterial genome catalog by Zhu et al^9^ were included, but genomes not isolated from oral cavity were manually identified and removed from the dataset.

### Metagenome-assembled genome (MAG) assembly

For the collected WMS data, adapter sequences were trimmed, and low-quality reads were removed using Trimmomatic (v0.39)^73^. The filtered reads were aligned to human genome (vCRCh38^74^) using Bowtie2 (v2.3.5.1)^75^ with the “--very-sensitive” option to remove contaminant human DNA sequence reads. Most processed samples were assembled into scaffolds using metaSPAdes (v3.15.3)^76^. For single-end sequencing samples and samples where metaSPAdes failed to complete assembly for unknown reasons, MEGAHIT (v1.2.9)^77^ was used instead.

Assembled scaffolds were binned to genome bins using three binning tools, MetaBAT2 (v2.12.1)^78^, MaxBin2 (v2.2.7)^79^, and CONCOCT (v1.1.0)^80^. To estimate read coverage and depth for scaffolds, processed reads were aligned back to the assembled contigs using Bowtie2 with the “--very-sensitive-local” option. Binning with MetaBAT2 was performed with a minimum contig size of 1,500, while MaxBin2 and CONCOCT were used with a minimum contig size of 1,100. Bin refinement was carried out using the result of all three binning methods as input for the bin_refinment module of MetaWRAP (v1.3.2)^81^.

### 16S rRNA sequence prediction from bacterial genomes

To identify 16S rRNA sequences from bacterial genomes, we employed Barrnap (v0.9)^82^ with an e-value threshold of 1e-5, resulting in the identification of 16S sequences in 1,802 genomes. Predicting 16S sequences is particularly challenging due to the difficulty of assembling 16S regions from short-read sequence data. For species where 16S rRNA sequences could not be predicted, we expanded our search to genome sequences from the same species in the GTDB r207^83^ database. This approach enabled the prediction of 16S rRNA sequences for an additional 670 species. As a result, we successfully identified 16S rRNA sequences for 2,161 out of 3,426 species (63%) in the HROM catalog.

### Evaluating genome quality and filtering chimeric genome

Genome completeness and contamination were initially assessed using the “lineage_wf” module of CheckM (v1.1.9)^19^. Additionally, tRNA sequences were identified and annotated using itRNAscan-SE 2 (v2.0.9)^84^ with the “-B” option for bacterial genomes and the “-A” option for archaeal genomes. For rRNA annotation, Barrnap (v0.9)^82^ was employed to identify 5S, 16S, and 23S rRNA sequences, applying e-value threshold of 1e-5 for 16S and 23S rRNA and 1e-4 for 5S rRNA. Genomes were categorized into three quality tiers based on specific criteria^85^. Medium-quality (MQ) genomes were defined as those with completeness ≥ 50%, contamination < 5%, and a genome quality score (Completeness – 5 X Contamination) ≥ 50. Near-complete (NC) genomes had a completeness ≥ 90%, contamination < 5%, and a genome quality score ≥ 50. High-quality (HQ) genomes met the same criteria as NC genomes but also contained 5S, 16S, and 23S rRNA sequences along with more than 18 tRNAs. To identify and exclude chimeric genomes, all genomes were analyzed with GUNC (v1.0.4)^22^. Chimeric genomes were defined as those with a clade separation score (CSS) > 0.45 and were removed from further analysis. Taxonomic classification of the remaining non-chimeric genomes was performed using the “gtdbtk classify_wf” module of GTDB-Tk2 (v2.1.1)^23^.

### Re-estimation of genome completeness

To address potential underestimation of genome completeness by the original CheckM^19^ “lineage_wf” workflow, which uses lineage-specific marker genes, we compared completeness values from “lineage_wf” with those from CheckM “taxonomy_wf” for the domain “Bacteria,” which relies on universal bacterial marker genes. Starting at the phylum level, we identified taxonomic clades with significantly lower completeness in the “lineage_wf” results using the Wilcoxon paired rank-sum test. We then focused on clades where genome quality classifications improved based on re-estimated completeness. For the underestimated clades, genome completeness was re-estimated using universal bacterial marker gene sets. In the case of the phylum Patescibacteria, which lacks many universal bacterial marker genes, we used CPR-specific marker genes^20^ for re-estimation. The newly estimated completeness values were used for the final genome quality classification. To validate the re-estimated completeness, we applied CheckM2^21^, a recently updated method that incorporated machine learning to address limitations of earlier workflows. We assessed the correlation between completeness values from CheckM “lineage_wf” and CheckM2, as well as the proportion of hits from the 120 single-copy marker genes used in GTDB-Tk, using Spearman correlation analysis.

### Species-level clustering and removal of redundant genomes

For genomes classified within the same order, MinHash distances were calculated using MASH^86^ (MASH sketch = 10,000), implemented in the dRep (v3.3.0)^87^ compare module with the options “--MASH_sketch 10000 --P_ani 0.8 --SkipSecondary”. Primary clusters were formed through hierarchical clustering with a distance threshold of 0.2. Within each cluster, the average nucleotide identity (ANI) was calculated for every genome pair using MUMmer4 (v4.0.0rc1)^88^. Genomes were then grouped into species clusters using average-linkage hierarchical clustering, implemented in R with the fastcluster pachage^89^. Clustering was based on ANI distance with a species-level cutoff of 95% identity (ANI distance of 0.05) and an alignment coverage threshold of 30% (0.3). For each species cluster, the representative genome was selected as the genome with the highest intactness score (*S*), calculated as *S* = *Completeness − 5 × Contamination + 0.5 × log_10_(N50)*. This scoring method follows the approach previously used in cataloging human gut microbial genomes^7^. To further refine the dataset, redundant genomes within species clusters were removed using average-linkage hierarchical clustering, applying an ANI cutoff of 99.9% identity (ANI distance of 0.001) and an alignment coverage cutoff of 81% (0.81). This process identified and removed 9,738 redundant genomes. Finally, a phylogenetic tree of the representative genomes was constructed using IQ-TREE2 (v2.1.3)^90^, based on the multiple sequence alignment of the 120 concatenated marker genes provided by GTDB-Tk (v2).

### Evaluation of sequence read classification performance

To assess the read classification performance of genome catalogs, including HROM, we obtained six additional WMS datasets from public repositories that were not used in the construction of HROM. Each dataset contains at least 30 samples, originating from various oral biogeographic sources. The datasets include buccal swab samples (PRJNA917836, n = 317), in-vitro oral biofilm samples (PRJNA983519, n = 75), oral samples (PRJNA983219, n = 30), saliva samples (PRJNA917836, n = 27; PRJNA997379, n = 45), supragingival plaque samples (PRJEB54673, n = 554), and tongue samples (PRJNA879058, n = 81), resulting in a total of 1,129 samples. Using these validation data, we tested the performance of several human oral microbial genome catalogs: eHOMD v3^10^, the Human Oral Genome Database by Zhu et al^9^, the Cultivated Oral Bacteria Genome Reference (COGR)^24^, and the human-oral-v.1.0.1 database from the MGnify repository^25^. For databases other than eHOMD, we applied the identical species clustering method and genome filtering scheme as used in HROM to build the near-complete species-level reference database built with representative genomes for Kraken2. All samples were aligned to each database using Kraken2^26^ with a confidence score threshold of 0.2 to assess their read classification performance.

### Constructing HROM custom database for marker-based species profiling

Marker-based species profiling was conducted using MetaPhlAn4^27^ with a custom database derived from HROM genomes. To construct database of species-specific marker genes, we adapted the MetaPhlAn4 computational pipeline. Panaroo (v1.3.0)^91^ was applied to 72,641 NC genomes and 3,246 species using the following options: “-c 0.90 --core_threshold 0.90 -f 0.5 -- clean-mode strict --merge_paralogs”, generating species-level pangenomes. For singleton species, genes/proteins from their representative species were used. Coding genes between 150–1,500 amino acids in length were selected from each species pangenomes and subsequently clustered using the linclust module of MMseqs2 (v2.1.9)^33^ with a 90% identity. To identify clade-specific, non-spurious markers, initial candidates with a coreness (prevalence among conspecific genomes) greater than 50% were selected, with the threshold increased to 60% for species with more than 100 conspecific genomes. To avoid duplicate genes or paralogs, markers with a mean count exceeding 1.5 per genome within a species were excluded. Additionally, markers that clustered with genes from more than 10 other species were discarded. The remaining marker candidates were fragmented into 150-nt segments and aligned to other species and conspecific genomes using Bowtie2. Markers aligned to more than 50% of conspecific genomes but fewer than 1% of genomes from any other species were retained. Only perfectly unique markers (aligning to no other species) or quasi-markers (aligning to fewer than 1% of genomes from other species) were included. After these filtering steps, up to 200 markers were selected per species to complete the custom marker database.

### Benchmarking species profiling methods

To evaluate taxonomic profiling methods using custom databases, we generated 5Gbp simulation datasets with CAMISIM (v1.3)^92^. From the 72,641 NC species in HROM, we randomly selected sets of 40, 150, 600, 1,000, and 1,500 species to simulate varying levels of species complexity within microbial communities. Simulations were performed using the CAMISIM “metagenomesimulation.py” module with the following configuration: “readsim = art_illumina, profile = mbarc, size=5”. For each genome set size, we generated 75 random samples under two strain complexity scenarios: single strain (selecting one genome per species) and multiple strains (selecting up to three genomes per species).

Taxonomic abundances were determined using both DNA-based and marker-based approaches. For the DNA-based approach, we employed Kraken2 paired with Bracken^28^ for species-level reassignment. Custom Kraken2 databases were generated using either representative genomes or concatenated conspecific genomes from HROM. Simulated reads were classified using Kraken2 with the options “--confidence = 0.2, --paired” for the respective HROM database. To minimize potential biases in DNA-based taxonomic abundance profiling, abundances were normalized by multiplying by the factor 10^6^ /*l_species_,* where *l_species_* is the mean genome length of the species^9^. Relative abundances were calculated by dividing normalized read counts by their total sum.

For marker-based profiling, microbiome profiles of simulated reads were estimated using MetaPhlAn4^27^ with the HROM custom marker database.

Performance metrics, including precision, recall, and F1-score, were calculated using Scikit-learn^93^ (v0.24.2). Bray-Curtis distances between ground-truth profiles and estimated profiles were computed using SciPy^94^ (v1.5.3). Hits to conspecific genomes within each species were treated as hits to the respective species.

Finally, to estimate the species count for each sample source from healthy individuals, we analyzed WMS samples with a sequencing depth greater than 5 Gbp using a custom pipeline and the HROM concatenated database. Species counts were derived from the species-level profiles generated for each sample.

### Construction of human reference oral microbial protein catalog

Protein coding sequence and its corresponding amino acid sequences were predicted for each HROM genome using Prokka (v1.14.6)^32^. The predicted amino acid sequences were concatenated and clustered at identity thresholds of 100%, 95%, 90%, 70%, and 50% using MMseqs2^33^ with the options “--kmer-per-seq 80 --cluster-mode 2”. This process generated protein catalogs for each identity threshold. Functional annotation of the protein catalogs was performed using eggNOG-mapper v2 (v2.1.9)^34,95,96^.

### Deep learning-based gene ontology annotation for microbial proteins

DeepGOPlus^37^(v1.0.1), a deep learning-based tool for gene ontology (GO) prediction, was used to annotate representative proteins from the protein clusters in the HROM dataset. Only predictions with a confidence score above 0.3 were retained. The GO term “cellular anatomical entity (GO:0110165)”, representing structural components within a cell, was excluded as it annotated 87.5% of the protein clusters, leading to an inflated prediction rate. Since DeepGOPlus was trained on data that includes eukaryotic protein sequences, some predictions might reflect functions specific to eukaryotes, potentially resulting in mispredictions for bacterial proteins. To address this, only bacteria-associated GO terms were retained. Proteins derived from bacteria were identified using the UniRef90 database (downloaded in September 2023), and their corresponding GO terms were designated as bacteria associated.

### Protein functional landscape analysis

For the protein cluster analysis, which included protein presence/absence and clustering of HROM with other protein catalogs, protein sequences inferred from representative genomes were clustered using MMseqs2 with the options “--min-seq-id 0.5 --kmer-per-seq 80 --cluster-mode 2”. The resulting presence/absence matrix, based on the clustering results, was imported as a table artifact into QIIME2 (v2022.11)^97^. The “q2-umap”^98^ module of QIIME2 was utilized to perform Uniform Manifold Approximation and Projection (UMAP) analysis, using the Jaccard distance matrix derived from protein content presence/absence data. Visualization of the UMAP results was conducted with ggplot2^99^.

To estimate the percentage of proteins clustered per species with the human gut microbiome, protein sequences from HROM representative genomes were clustered with the protein catalog of HRGM2 using MMseqs2 with the same option. For each HROM representative genome, the number of proteins clustered with HRGM2 proteins was identified, and the ratio of shared proteins was calculated by dividing the number of clustered proteins by the total protein count of the species. For clustering percentage with proGenomes3^31^, the representative protein sequence of proGenomes3 were obtained, and the same process was applied to infer clustering results.

### Analysis of species-level pangenome and single nucleotide variants (SNVs)

For species-level pangenome analysis, we conducted Panaroo (v1.3.0)^91^ with the options of “-c 0.90 --core_threshold 0.90 -f 0.5 --clean-mode strict --merge_paralogs”, applied to GFF files produced by Prokka for NC conspecific genomes per species. From the pangenome gene presence/absence matrix, we generated the rarefaction curves by randomly removing genomes.

The resulting curve was fitted to the equation y ∼ log(x), where *x* is the number of genome and *y* is the size of the pangenome. For the analysis of single nucleotide variants (SNVs) in HROM, we utilized scripts and pipeline provided in previous research^6^ (available at https://github.com/zjshi/snv_analysis_almeida2019). For species with more than five conspecific genomes, each non-representative conspecific genome (*g*) was aligned against the representative genome (*r*) of the species cluster using nucmer from MUMmer4^88^. Best bi-directional alignments were identified using the delta-filter program with options “-q –r”. SNVs were annotated with the show-snp program of MUMmer4. For each species with a representative genome (*r*), the SNV density was calculated by normalizing the total number of SNVs by the aligned genome length and the number of conspecific genomes (*G*) as follows:

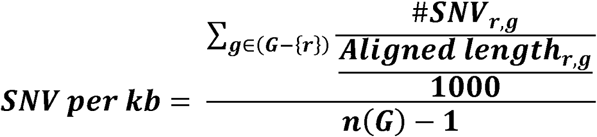

For downstream analyses and SNV density calculations, only species with more than 10 NC conspecific genomes were included.

### Identifying biosynthetic gene clusters (BGCs) and metabolic gene clusters (MGCs)

Biosynthetic and metabolic gene clusters were identified for all NC genomes in HROM. MGCs were detected using gutSMASH (v1.0.0)^100^,while secondary metabolite BGCs were identified using antiSMASH (v6.1.1)^101^.

### Profiling division and cell wall (dcw) gene clusters

To profile division and cell wall (dcw) gene clusters, we obtained 17 dcw gene cluster HMM models from the original research dataset (available at https://doi.org/10.17632/4y5mzppzmb.1).

The HMM models were used with hmmsearch (e-value threshold = 1e-5) to identify hits for each marker within the genomes.

### Profiling genomes for antibiotic resistance and viral defense systems

Antibiotic resistance genes in the 72,641 NC genomes of HROM were identified using the “RGI main” function (with the “strict” option) of RGI (v6.0.3)^102^. Viral defense systems were inferred using DefenseFinder (v1.0.9)^103^.

### Identifying anti-microbial peptides (AMPs)

To identify antimicrobial peptides (AMPs), amino acid sequence of small open reading frame (smORF) were inferred from all NC genomes in HROM using Prodigal (v2.3.0) with the option “--min-gene = 33”^104^. The smORFs were filtered to include sequences raging between 15 and 100 amino acids. To remove spurious smORFs, we employed the AntiFam^105^ HMM model and performed sequence filtering using hmmsearch. Redundant smORFs were then removed to generate a non-redundant dataset. The non-redundant smORFs were subject to AMP prediction using Macrel (v1.2.0)^39^. Singleton smORFs identified as AMP were excluded. To estimate the proportion of novel AMPs among those derived from HROM genomes, we obtained reference AMPs from the database of Antimicrobial Peptides (DRAMP v3.0)^18^, ensuring that the sequences were correctly labeled with amino acids. AMPs derived from HROM were clustered with AMPs retrieved from DRAMP at the family-level using cd-hit of CD-HIT (v4.6.8)^106^ with the options “-c 0.75 -M 0 -d 0” after converting the amino acids sequences into a reduced alphabet of eight letters^107^. The AMP density for each genome was defined as number of AMPs identified divided by the total genome length.

### Patescibacteria genomes from different isolation sources

We collected genomes annotated as Patescibacteria from GenBank. Additionally, we collected those isolated from groundwater ecosystems^54^. These genomes were classified using GTDB-Tk^23^ and only those confirmed to belong to the phylum Patescibacteria were retained. The genomes were subjected to the same quality filtering criteria used for HROM to ensure only non-chimeric, NC genomes were included. Metadata were manually inspected to determine the isolation source of each genome. Genomes were annotated as MHA (mammalian host-associated) if isolated from animal or human associated environments; otherwise, they were categorized as “environment”. Using the HROM representative genomes classified as Patescibacteria, we applied the same species-level de-replication pipeline as HROM to infer species-level genome bins within Patescibacteria. For protein family analysis, protein sequences were predicted from the representative genome of each species using Prokka^32^. These sequences were then clustered and analyzed using the same methodology described earlier to infer the protein family of Patescibacteria.

### Inference of protein family from Patescibacteria genomes

Protein families were inferred using a pipeline and script from a previous study (available at https://github.com/raphael-upmc/proteinClusteringPipeline)^47^. Briefly, an all-vs-all similarity search was conducted using MMseqs2^33^ with the following parameters: e-value = 0.001, sensitivity = 7.5, and coverage = 0.5. The input consisted of protein sequences derived from the representative genome of 2,963 Patescibacteria genomes. A similarity-based network was constructed and used as input for clustering protein sequences into subfamilies using the greedy set cover algorithm of MMseqs2. Subfamilies containing more than two protein members were selected for downstream analysis. These selected subfamilies were aligned using the “mmseqs result2msa” module of MMseqs2, and multiple sequence alignments were used to construct HMM profiles with the HHpred suite^108,109^. Pairwise comparisons of the HMM profiles were performed using hhblits from the HHpred suite with the following parameters: -v 0 -p 50 -z 4 -Z 32000 -B 0 -b 0. Subfamilies with probability scores ≥95% and coverage ≥0.50 were integrated into the input network, with similarity scores serving as edge weights. Protein families were defined by clustering the network using the Markov Clustering (MCL) algorithm with an inflation parameter of 2.0.

### Functional enrichment analysis and metabolic comparison of Patescibacteria genomes

For functional term enrichment and metabolic comparison, we profiled each representative genomes of 2,963 Patescibacteria species using the “anvi-run-ncbi-cogs” and “anvi-run-kegg-kofams” modules of Anvi’o. Additional functional terms were identified using MacSyFinder (v2)^110^ and InterProScan (v5.60-92.0)^111^. The completeness of KEGG metabolism modules was assessed using “anvi-estimate-metabolism” module, with modules considered present if their completeness was equal to or greater than the default threshold of 0.75. Terms and proteins were defined as prevalent if their prevalence among MHA genomes was ≥ 50% and their prevalence among environmental genomes was ≤ 20%.

### Analyses of the periodontitis-associated Patescibacteria

For species clustering based on protein profiles, we used the “makeSNNGraph” function from the R package bluster (v1.9.1)^112^ to construct a nearest-neighbor graph. Louvain clustering was performed using the “cluster_louvain” function from the R package igraph (v1.3.1)^113^. Four public datasets, comprising both healthy and periodontitis samples, were included in the analysis. Differentially enriched Patescibacteria clusters were identified using MaAsLin2 (v1,8)^114^, with the batch variable treated as a random effect. Batch correction for the saliva periodontitis datasets was performed using MMUPHin (v1.16)^115^.

To identify protein families enriched in periodontitis-associated Patescibacteria cluster 16, we selected families with a prevalence lower than 5% in other clusters but higher than 50% in cluster 16. WMS reads were realigned to the selected protein families using DIAMOND (v2.1.9)^96^ with threshold of 80% coverage and 80% identity. Any aligned reads to these proteins were considered hit. The relative abundance of protein families was estimated by dividing the number of hits by the total number of reads.

For protein structure prediction, we utilized ESMFold (v1.0.3)^60^, followed by a protein structure search using Foldseek (v9.427df8a)^62^ against the Alphafold UniProt50^61^ database, employing the “easy-search” mode. Hits were retrieved if they met the following criteria: an e-value less than 10-4, a TM score ≥ 0.3, and query coverage ≥ 0.5.

### Identification of oral-gut shared species

Species-level de-replication was performed for representative genomes from HROM (oral) and HRGM2 (gut) using the previously described pipeline. In brief, all representative genomes were taxonomically classified with GTDB-Tk^23^. For genomes within the same order, primary clusters were defined using the MASH tool from the dRep compare module, applying the same parameters as those used in HROM species definition. Within each primary cluster, ANI was calculated, and hierarchical clustering was conducted using the same parameter to achieve species-level de-replication. Genome bins that included at least one genome from HRGM2 and one from HROM were identified as oral-gut shared species.

### Estimation of sporulation capacity

To quantify the sporulation score, we followed a scheme similar to that used in a previous study^70^. From the 66 known sporulation-predictive genes, we successfully retrieved protein sequences for 65 of them. The presence of 65 sporulation-predictive genes in each genome was measured using tblastn from BLAST (v2.12.0+)^116^ with the parameters e-value = 1e-5 and 30% identity. The sporulation score was calculated as the number of sporulation-predictive genes detected in the genome divided by 65. For each species, the highest sporulation score observed among its conspecific genomes was used as the representative sporulation score.

### Estimation of metabolic independence

To estimate metabolic independence, we followed a scheme similar to that described in a previous study^69^. The metabolic independence score was calculated as the sum of the completeness scores of 33 KEGG modules associated with metabolic independence. KEGG modules were identified using anvi-run-kegg-kofams function from Anvi’o (v7.1)^117^. The completeness of each KEGG metabolism module was then quantified using the anvi-estimate-metabolism module of Anvi’o, with a default threshold of 0.75 applied to confirm module presence. The metabolic independence score was defined as total completeness scores of the 33 KEGG modules divided by 33. For each species, the representative metabolic independence score was calculated as the mean score of all its conspecific genomes.

### Identification of ectopic oral species

To identify ectopic oral species, which are presumed to be prevalent in the healthy human oral cavity but not in the healthy gut microbiota, we analyzed 15 healthy oral microbiome datasets (**Supplementary Table 15**) and 9 healthy gut microbiome datasets (**Supplementary Table 16**), each containing at least 30 samples. For oral microbiome datasets, if a single dataset included samples from more than two isolation sources, each subset was treated as a separate dataset. This resulted in the collection of 3,377 healthy oral samples and 956 healthy gut samples for analysis. For the healthy oral microbiome datasets, species-level classification of WMS reads was conducted using the HROM custom database based on concatenated genomes. Similarly, for healthy gut microbiome datasets, species-level classification was performed using the HRGM2 custom database. From these species-level profiles, we identified 330 oral-gut shared species and measured their prevalence using a relative abundance threshold of 1e-6 to determine species presence. For cases where a single HROM species was clustered with multiple HRGM genomes, the highest prevalence value of the HRGM2 species was selected. Conversely, when multiple HROM species were clustered with a single HRGM2 genome, the prevalence of the HRGM2 genome was assigned to all clustered species. The mean prevalence was calculated across all datasets for each species within oral and gut samples. Thresholds of 50% prevalence in healthy oral microbiomes and 25% prevalence in healthy gut microbiomes were applied to define the ectopic oral species.

### Identifying important bacterial species for colorectal cancer (CRC) classification

We analyzed 12 cohorts of human gut microbiome samples from colorectal cancer (CRC) patients and healthy controls (**Supplementary Tables 18–19**). These datasets were processed using a workflow similar to the preprocessing of WMS data for HROM. Taxonomic classification was performed with Kraken2 using the HRGM2 custom genome database. Species-level counts were estimated using Bracken, and reads were normalized by multiplying by 10^6^/*l_species_*, where *l_species_* represents the mean genome length of the species. Relative abundance was calculated by dividing the normalized abundance by their total sum. From the multi-cohort profile, we identified differentially abundant features associated with CRC using the “check.associations” function (option = “lm”, formula = “feat ∼ cohort + disease”) of SIAMCAT (v2.6.0)^118^, applying an FDR-adjusted *p*-value ≤ 0.05. Additionally, we employed ANCOM-BC^119^ in individual cohorts to identify CRC-enriched taxa with the same FDR-adjusted *p*-value threshold. The variance explained by cohort and disease labels was assessed using “check.confounders” function of SIAMCAT. For CRC classification, we trained a random forest model on gut species profiles from nine datasets using SIAMCAT functions “create.data.split” and “train.model” (option = “method = randomForest”). CRC prediction was then conducted on the remaining three datasets not used for training, using the SIAMCAT functions “make.predictions” and “evaluate.predictions”. To identify the key features of the model, we used the “feature_weights” function to extract the median relative feature weights. The top 20 species with the highest median relative feature weights were designated as the most important features. To construct microbial networks, we employed SPIEC-EASI (v1.1.0)^120^. Specifically, we focused on 17 oral bacterial species among the top 20 important species for CRC classification.

## Data availability

By accessing the web server, www.decodebiome.org/HROM/ users can browse and download sequences and annotations for oral microbial genomes, protein families, 16S regions, a custom Kraken2 database, and a database of species-specific markers.

## Acknowledgements

This work was supported by the National Research Foundation (NRF) funded by the Ministry of Science and ICT, Republic of Korea (2022M3A9F3016364, 2022R1A2C1092062), Ministry of Trade, Industry and Energy (200022947), Brain Korea 21 (BK21) FOUR program.

## Author contributions

J.H.C. and I.L. conceived and designed the study. J.H.C. performed the construction of genome catalog and formulated the study hypothesis. N.K., J.M. co-developed bioinformatics pipelines. S.L., G.K. assisted data collection and compilation. Y.S. assisted building the data server. S.B., I.B., and B.L. assisted data analysis. I.L. supervised the project. J.H.C. and I.L. wrote and edited the manuscript.

## Declaration of generative AI and AI-assisted technologies in the writing process

During the preparation of this work the authors used ChatGPT in order to improve language and readability, with caution. After using this tool, the authors reviewed and edited the content as needed and take full responsibility for the content of the publication.

## Figures

**Extended Data Fig. 1.**
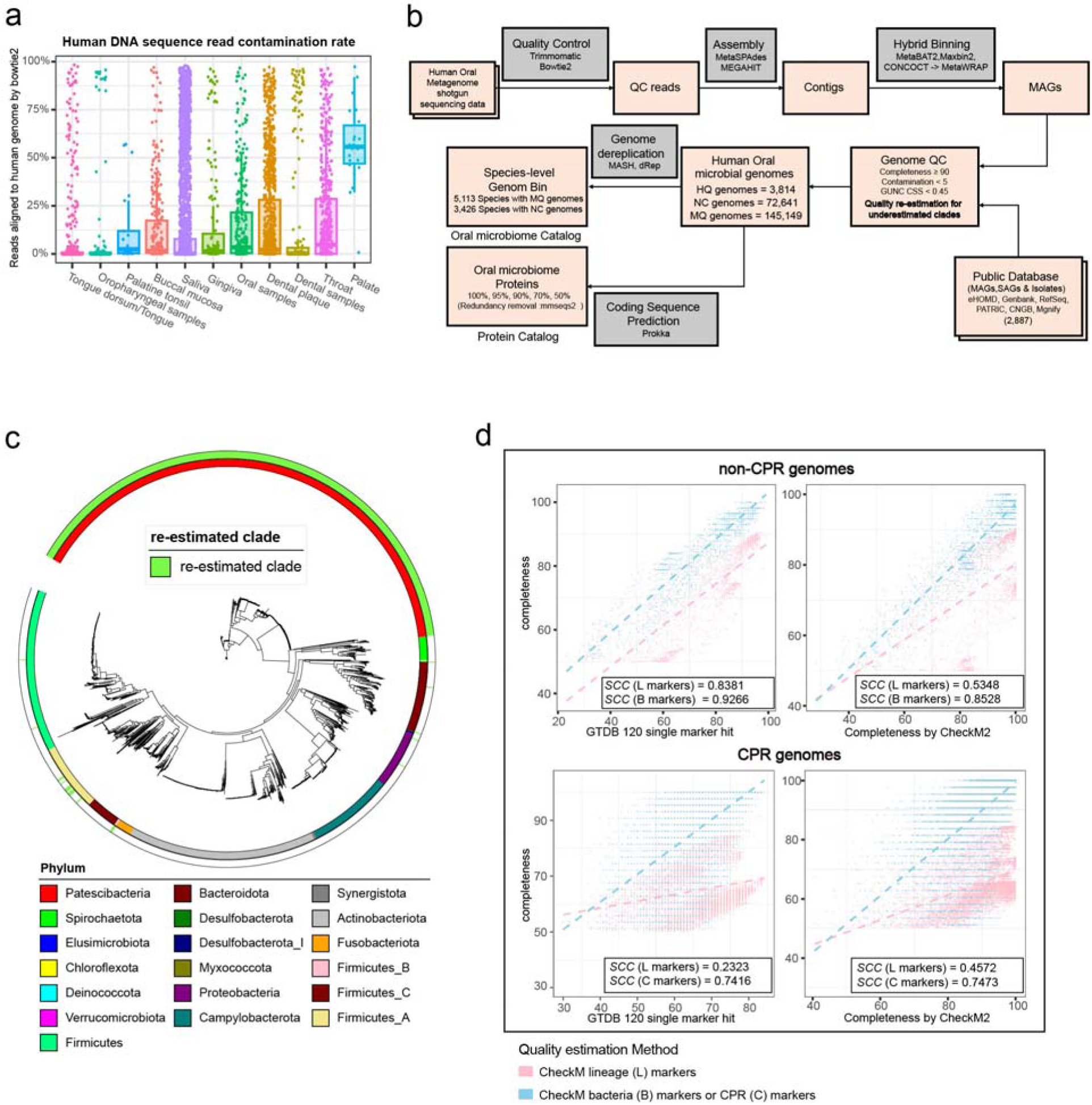
Quality control and completeness re-estimation of oral microbial genomes. **a**, Boxplot showing the human DNA sequence contamination rates of whole metagenome sequencing datasets across various oral sample types. Contamination rates were estimated by aligning reads to the human reference genome, GRCh38, using Bowtie2. Boxplot elements: center line (median), box edges (25th and 75th percentiles), whiskers (1.5× interquartile range). **b**, Bioinformatics workflow for the construction of the Human Reference Oral Microbiome (HROM) catalog, including quality control (QC), genome assembly, binning, and completeness re-estimation for underestimated clades. The pipeline integrates publicly available databases and hybrid binning methods to produce species-level genome bins and a comprehensive protein catalog. **c**, Phylogenetic tree highlighting clades with re-estimated genome completeness. Green bars indicate clades where completeness was re-estimated due to underestimation using lineage-specific markers. Taxonomic assignments are color-coded by phylum. Most of the re-estimated genomes belong to phylum Patescibacteria. **d**, Scatterplots comparing genome completeness estimates across different methods for non-CPR (top row) and CPR (bottom row) genomes. Completeness was evaluated using CheckM with lineage-specific markers (L markers), universal bacterial markers (B markers), and CPR-specific markers (C markers). Spearman correlation coefficients (SCC) between the methods and GTDB 120 single marker hits or CheckM2 completeness are shown for each plot.

**Extended Data Fig. 2.**
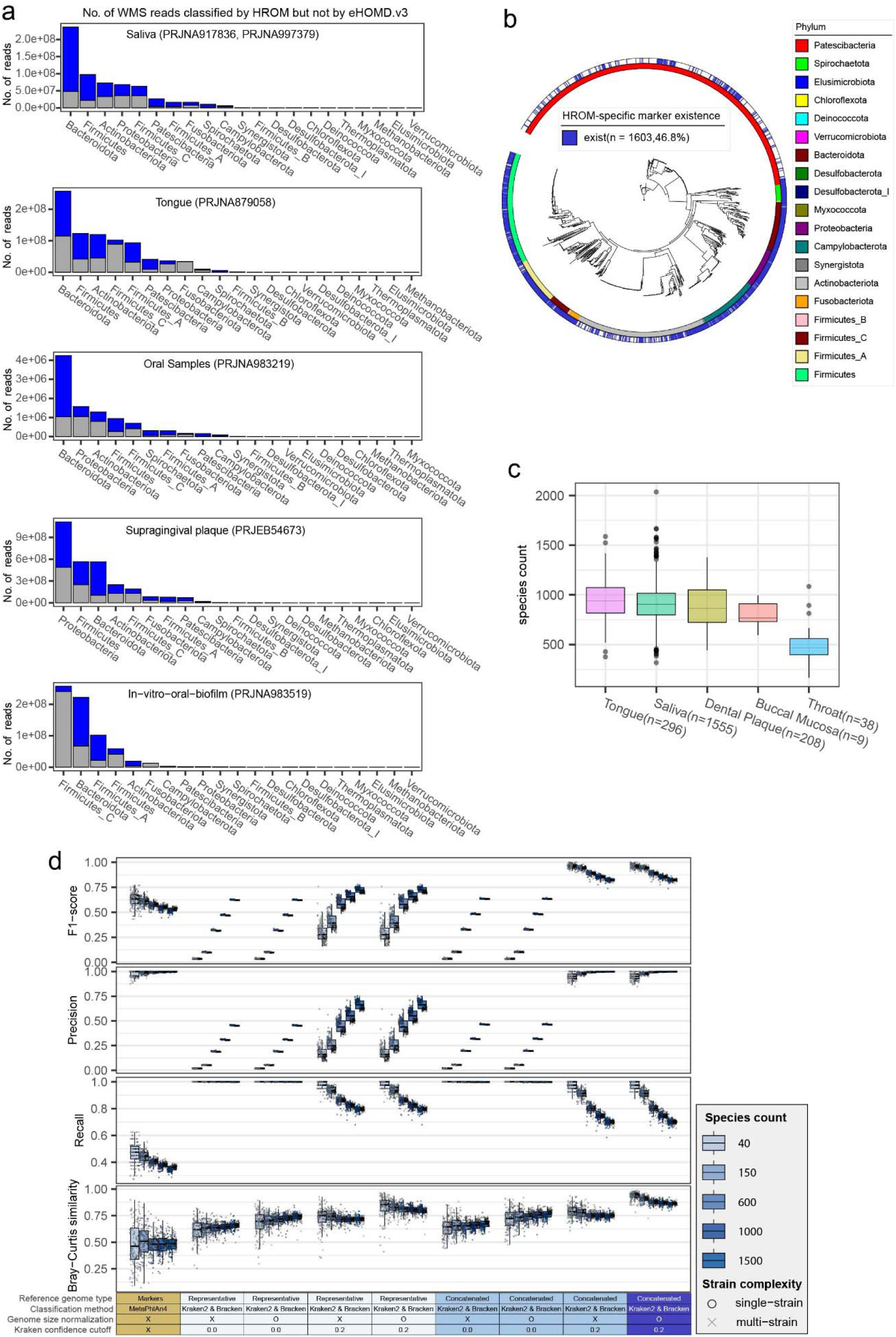
Benchmarking and expanded classification capability of the HROM catalog. **a**, Number of whole metagenome sequencing (WMS) reads classified by HROM but not by eHOMD v3 across various oral sample types, including saliva, tongue, oral samples, supragingival plaque, and in-vitro oral biofilm. Taxonomic breakdown highlights the significant contribution of HROM in classifying previously unclassified reads across diverse phyla. **b**, Phylogenetic tree of HROM representative genomes with tracks indicating the availability of species-specific marker genes (blue), with 46.8% of genomes showing species-specific marker existence. Phyla are color-coded. **c**, Boxplot of species counts per sample across different oral sample sources, including tongue, saliva, dental, buccal plaque, and throat mucosa. Sample sizes for each source are indicated. **d**, Benchmarking of species profiling performance across varying levels of species and strain complexity using different classification methods (MetaPhlAn4 with a custom HROM species-specific marker database, Kraken2/Bracken with a custom HROM representative genome database, and Kraken2/Bracken with a custom HROM concatenated genome database). The evaluation considered variations with and without genome length normalization and a confidence score cutoff 0.2. Metrics include F1-score, precision, recall, and Bray-Curtis similarity, highlighting the superior performance of HROM in both single-strain and multi-strain scenarios. (c-d) Boxplot elements: center line (median), box edges (25th and 75th percentiles), whiskers (1.5× interquartile range).

**Extended Data Fig. 3.**
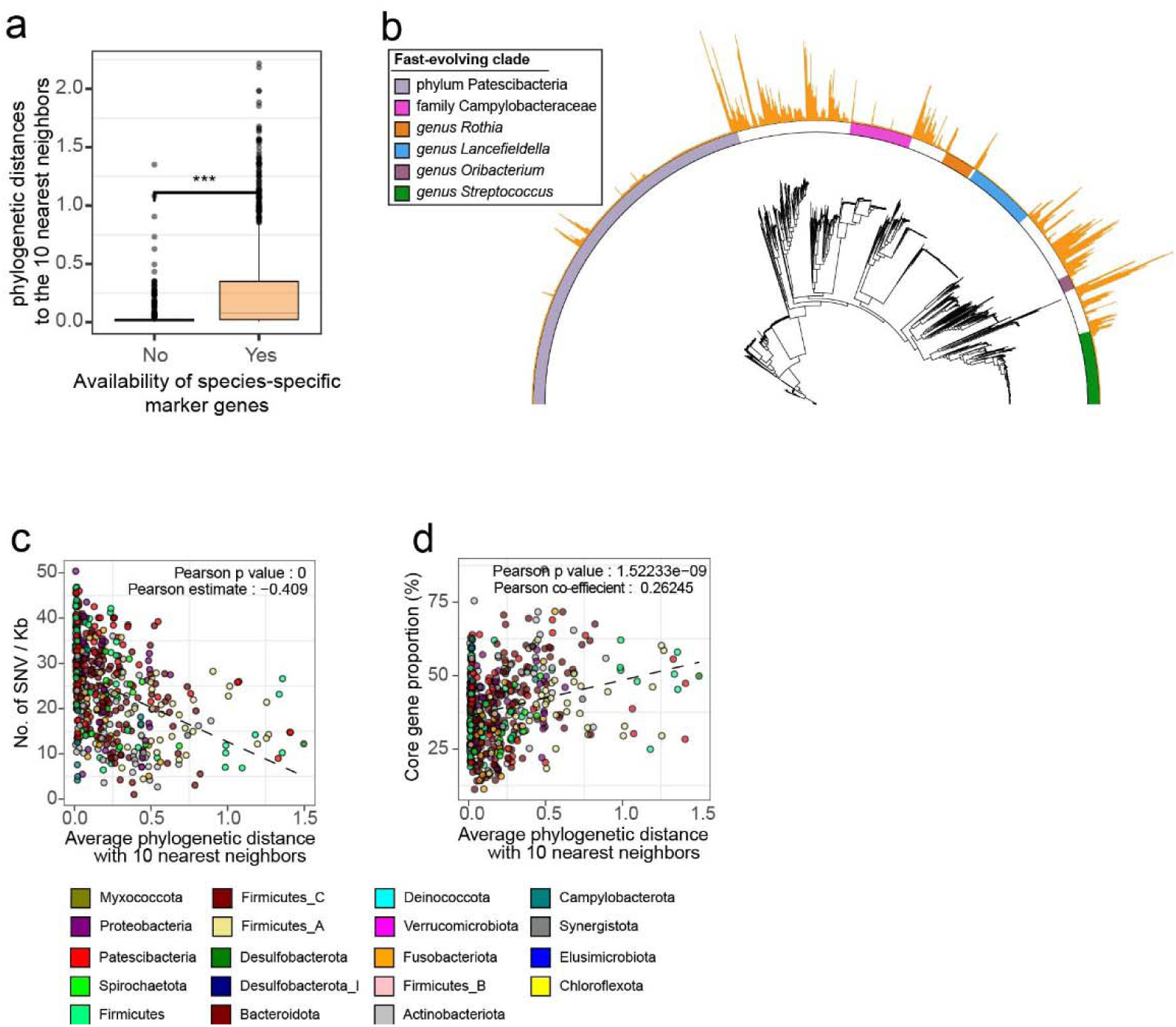
Phylogenetic and genomic features of fast-evolving oral microbial clades. **a**, Boxplot comparing the phylogenetic distances to the 10 nearest neighbors for species with (Yes) and without (No) species-specific marker genes. Species lacking specific markers exhibit smaller phylogenetic distances, indicating closely related neighbors. Boxplot elements: center line (median), box edges (25th and 75th percentiles), whiskers (1.5× interquartile range). Statistical significance was determined using a two-tailed Mann-Whitney U test (*** *P* < 0.001). **b**, Phylogenetic tree of oral microbiota highlighting six clades undergoing rapid speciation. Outer tracks indicate phylogenetic distances to the 10 nearest neighbors (orange bars) and taxonomic classification (color-coded by phylum). **c**, Scatterplot showing the relationship between single nucleotide variant (SNV) density (the number of SNVs per kb) and the average phylogenetic distance to the 10 nearest neighbors. **d**, Scatterplot showing the relationship between the proportion of core genes (%) and the average phylogenetic distance to the 10 nearest neighbors.

**Extended Data Fig. 4.**
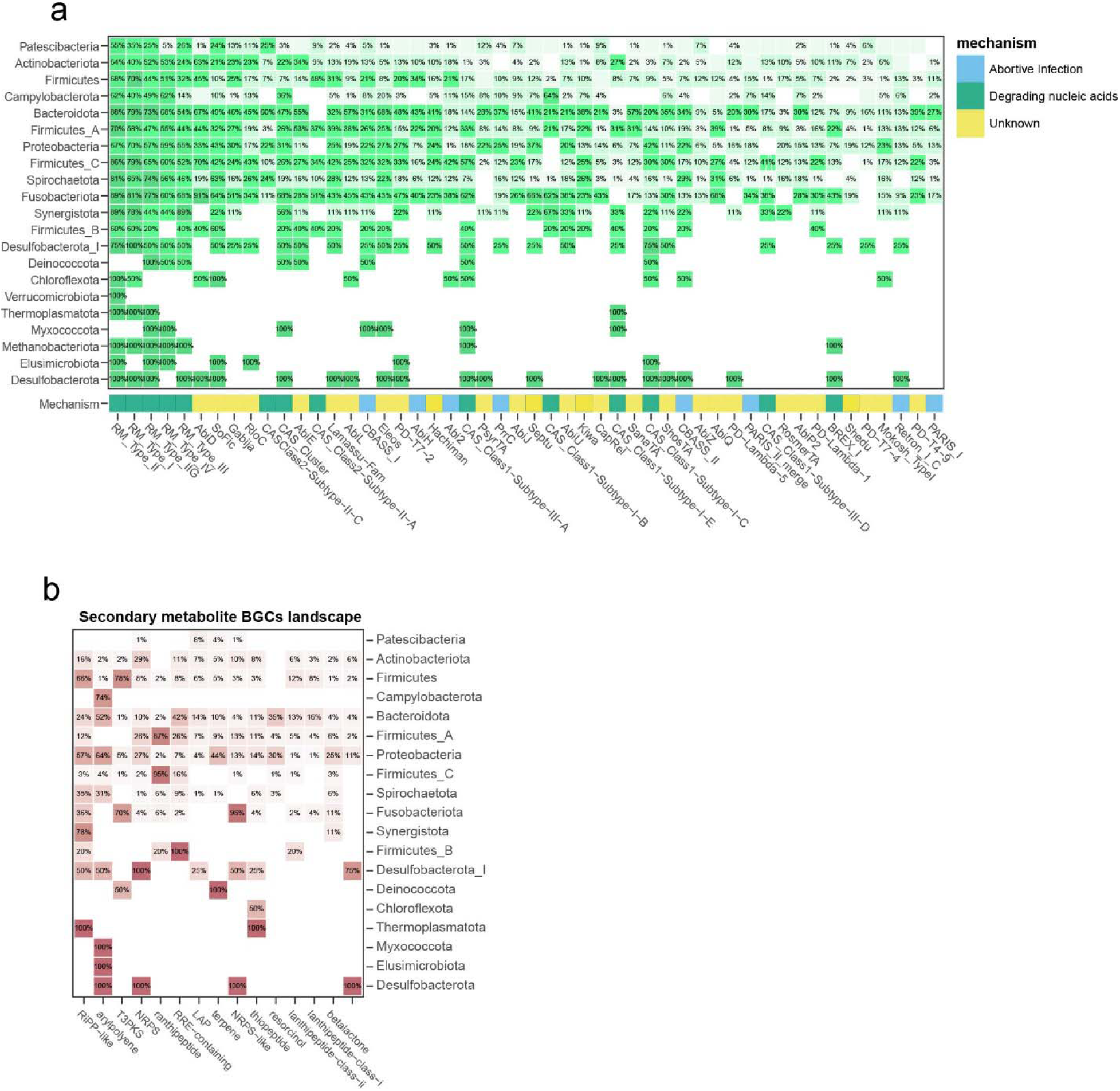
Functional landscape of viral defense mechanisms and secondary metabolite BGCs in oral microbial phyla. **a**, Heatmap showing the distribution of viral defense mechanisms across oral microbial phyla. Rows represent phyla, and columns correspond to specific viral defense mechanisms. Prevalence percentages are displayed for each phylum and mechanism. Viral defense mechanism was considered present if it was identified in at least one conspecific genome within the same species. **b**, Heatmap representing the distribution of secondary metabolite biosynthetic gene clusters (BGCs) across major oral microbial phyla. Rows correspond to phyla, and columns indicate specific types of BGCs. The prevalence of each BGC type within phyla is shown as percentages, highlighting the diversity of secondary metabolite production capabilities among phyla. BGCs was considered present if it was identified in at least one conspecific genome within the same species.

**Extended Data Figure 5.**
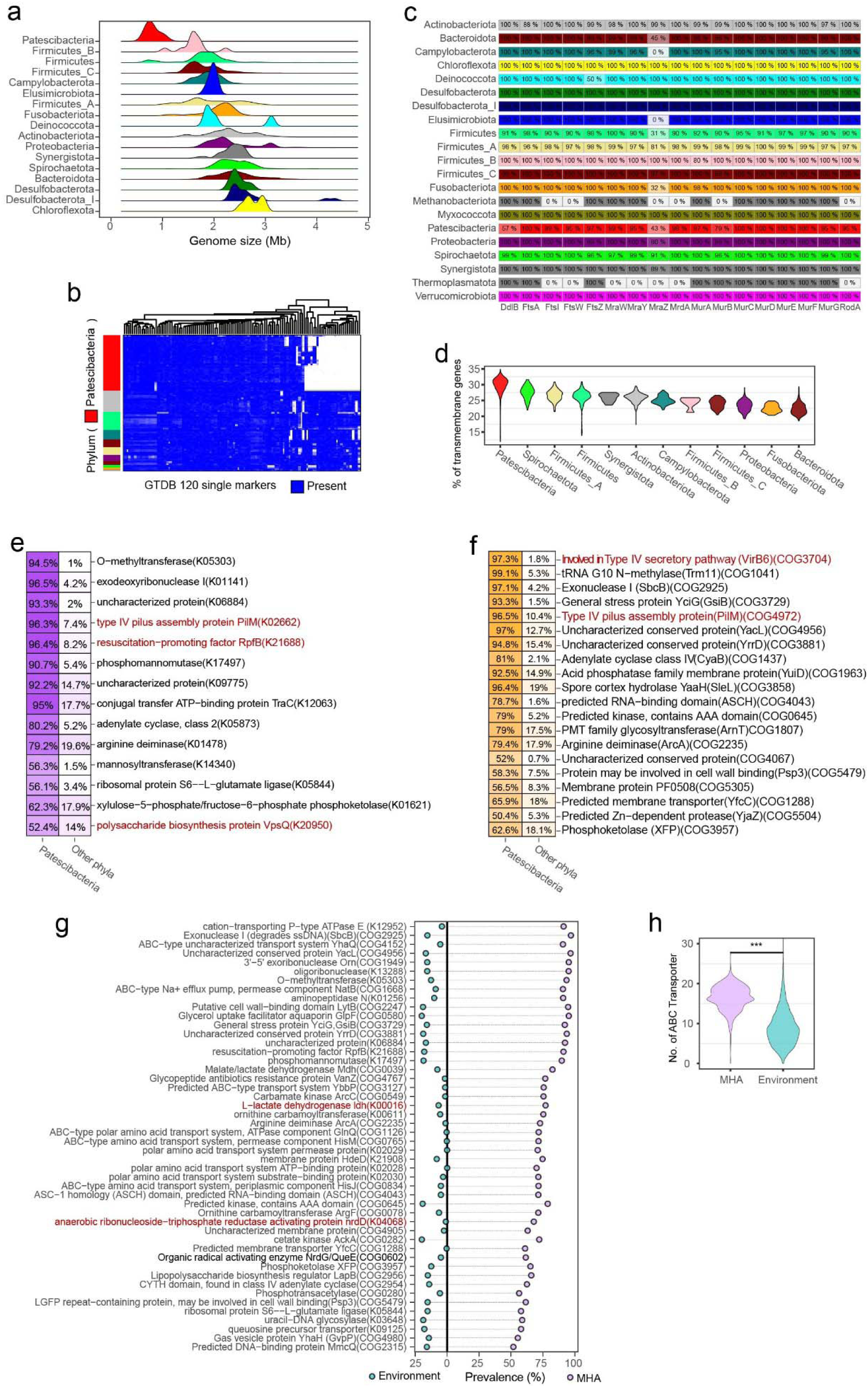
Comparative genomic and functional analyses of Patescibacteria. **a**, Ridge plot showing HROM representative genome size distribution across major phyla, with Patescibacteria exhibiting the smallest genome sizes. **b**, Heatmap of GTDB 120 single marker gene presence across 3,426 HROM representative species genomes. The left color strip indicates the phylum of each species, while the column represents the GTDB 120 single markers. A red strip identifies the phylum Patescibacteria. **c**, Heatmap showing the presence of division and cell wall (*dcw*) genes across phyla. Each row corresponds to a phylum, and columns represent specific *dcw* genes. The percentage inside each box denotes the prevalence of corresponding gene for each phylum. **d**, Violin plot displaying the proportion of transmembrane genes across phyla, with Patescibacteria showing higher distribution compared to other phyla. **e-f**, Table listing the top enriched KEGG orthologs (**e**) and COG terms (**f**) in Patescibacteria genomes compared to other phyla, with percentages indicating prevalence within Patescibacteria and the relative enrichment compared to other phyla. **g**, Lollipop chart showing the prevalence of functional terms in Patescibacteria genomes, comparing mammalian-host-associated (MHA) and environment species. Blue points indicate prevalence in environment species, while violet points indicate prevalence in MHA genomes. **h,** Violin plot comparing the number of ABC transporter protein in MHA versus environment Patescibacteria genomes, with MHA genomes showing significantly more transporters. Statistical significance was assessed using a two-tailed Mann-Whitney U test (*** *P* < 0.001).

**Extended Data Figure 6.**
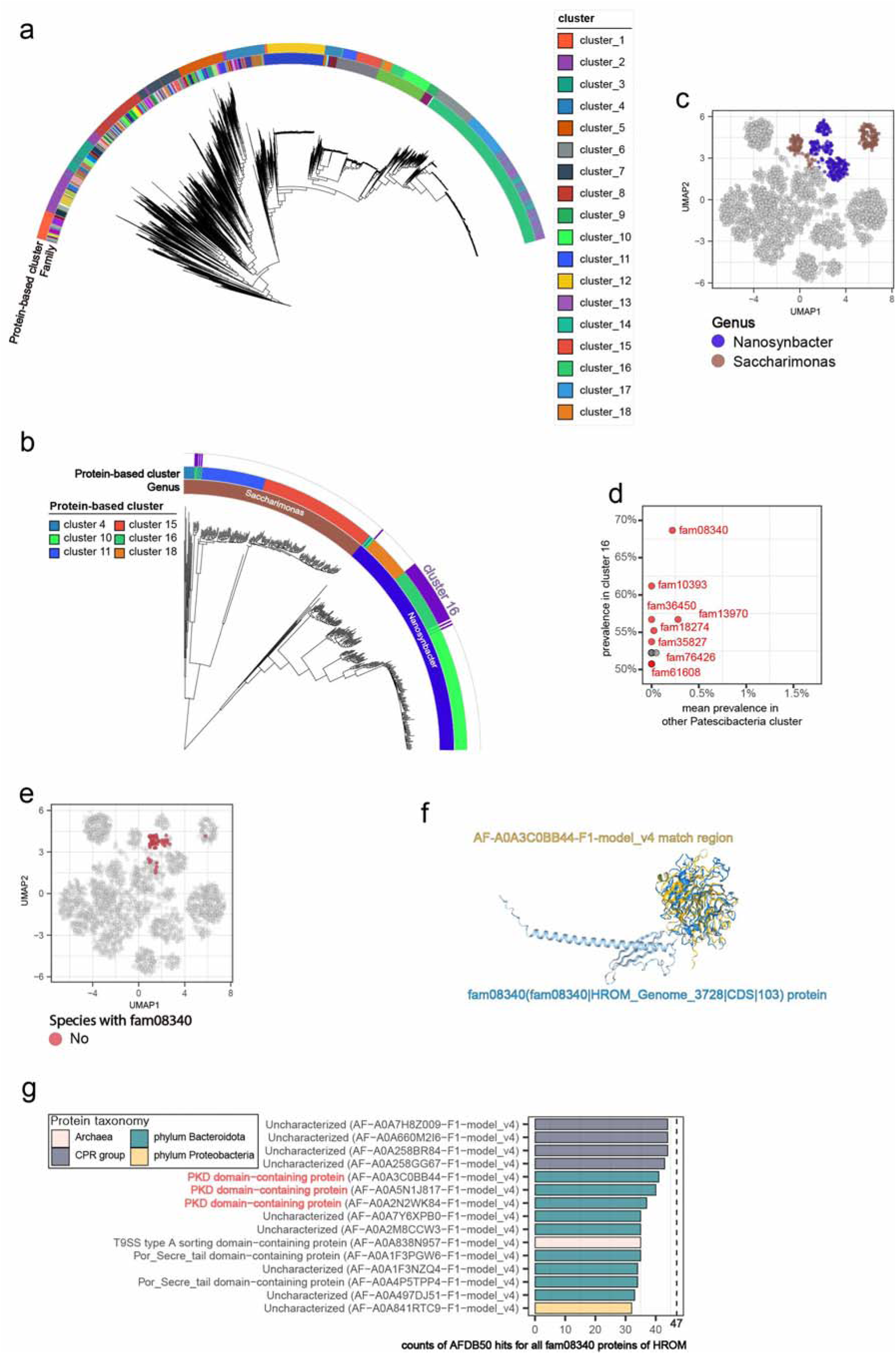
Phylogenetic and functional insights into Patescibacteria clusters. **a**, Phylogenetic tree highlighting clusters of 2,963 Patescibacteria species based on the occurrence profile of protein families at 50% sequence identity. The first strip indicates the Family, while the second strip represents clusters based on protein family presence/absence. **b**, Detailed phylogenetic view of 402 species from Patescibacteria clusters 4, 10, 11, 15, 16, and 18, with annotations indicating associated genera (*Saccharimonas* and *Nanosynbacter*). **c**, UMAP visualization of Patescibacteria species based on protein family profiles. Species belonging to the genera Nanosynbacter and Saccharimonas are color coded. **d**, Scatterplot displaying the prevalence of protein families in cluster 16 (y-axis) compared to its mean prevalence across other Patescibacteria clusters (x-axis). **e**, UMAP visualization showing species containing the enriched protein family fam08340 (red points). **f**, Predicted protein structure of a representative fam08340 protein using ESMFold, with highlighted functional match regions. **g**, Bar plot illustrating structural search hits using Foldseek. The bars represent the number of proteins with hits, while the vertical line indicates the total number of proteins in fam08340.

**Extended Data Fig. 7.**
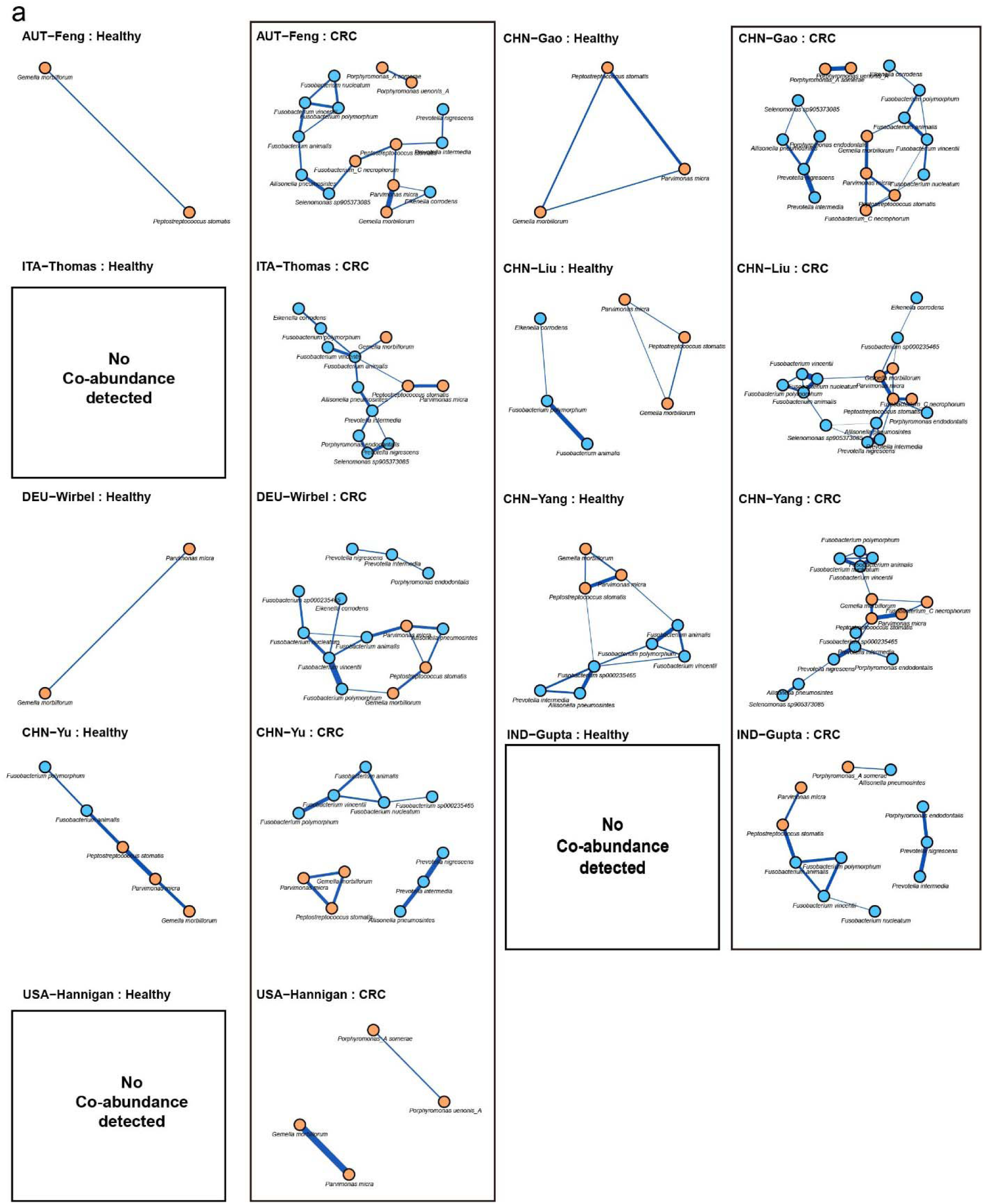
Co-abundance networks of 17 oral species among top 20 important bacterial species for colorectal cancer (CRC) classification. **a**, Each panel represents the co-abundance relationships among the 17 oral species including oral-gut shared (orange) and ectopic oral (blue) species from the top 20 important species for CRC classification. Edges indicate significant co-abundance relationships. Co-abundance was more frequently detected in CRC samples compared to healthy control samples, highlighting the altered microbiome interactions in CRC. Cohorts where no significant co-abundance relationships were detected in healthy samples are marked as “No Co-abundance Detected.” This pattern emphasizes the association of oral bacterial species with CRC-associated gut microbiome dysbiosis.

